# TAPBPR Promotes Antigen Loading on MHC-I Molecules Using a Peptide Trap

**DOI:** 10.1101/2020.04.24.059634

**Authors:** Andrew C. McShan, Christine A. Devlin, Giora I. Morozov, Sarah A. Overall, Danai Moschidi, Neha Akella, Erik Procko, Nikolaos G. Sgourakis

## Abstract

Chaperones tapasin and TAP-binding protein related (TAPBPR) perform the important functions of stabilizing nascent MHC-I molecules (chaperoning) and selecting high affinity peptides in the MHC-I groove (editing). While X-ray and cryo-EM snapshots of MHC-I in complex with TAPBPR and tapasin, respectively, have provided important insights into the peptide-deficient MHC-I groove structure, the molecular mechanism through which these chaperones influence the selection of specific amino acid sequences remains incompletely characterized. Of particular importance is a 16 residue loop in TAPBPR (corresponding to 11 residues in tapasin), which has been proposed to actively compete with incoming peptides by forming direct contacts in the F-pocket of the empty MHC-I groove. Using a deep mutational scanning functional analysis of TAPBPR, we find that important residues for the chaperoning activity are located on the major interaction surfaces with nascent MHC-I molecules, excluding the loop. However, interactions with properly conformed molecules toward peptide editing are influenced by loop mutations, in an MHC-I allele- and peptide-dependent manner. Detailed biophysical characterization by ITC, FP and NMR reveals that the loop does not interact with the peptide-deficient MHC-I groove to compete with incoming peptides, but instead promotes peptide loading by acting as a kinetic trap. Our results suggest that the longer loop of TAPBPR lowers the affinity threshold for peptide selection, to promote loading within subcellular compartments of reduced peptide concentration and to prevent disassembly of high affinity peptide-MHC-I complexes that are transiently interrogated by TAPBPR during editing.

## Introduction

Class I major histocompatibility complex (MHC-I) molecules bind a repertoire of endogenously processed peptide antigens and display them on the cell surface, thereby enabling CD8^+^ cytotoxic T lymphocytes to surveil the cell proteome (1). The disparate surface chemistries of peptide/MHC-I (pMHC-I) antigens encompassing self, pathogen or tumor derived peptides allows T lymphocytes to detect aberrant protein expression and mount a response toward infected or tumor cells (2). Assembly of pMHC-I molecules occurs in the endoplasmic reticulum (ER), where the association between a peptide, the polymorphic heavy chain (human leukocyte antigen, HLA), and the invariant light chain (β_2_-microglobulin, β_2_m) is facilitated by dedicated molecular chaperones, the ER-restricted tapasin of the peptide-loading complex (PLC) and the PLC-independent TAPBPR (TAP-binding protein related) (3, 4). In addition to chaperoning MHC-I, tapasin and TAPBPR function as catalytic enhancers of peptide association and dissociation within the MHC-I groove (peptide exchange) and participate in peptide editing and quality control of assembled pMHC-I molecules (5–8). Collectively, these processes ensure the correct folding, trafficking and prolonged cell surface display of pMHC-I antigens (9). Improper chaperone function is associated with severe impairment of the adaptive immune system and results in accelerated disease progression in the context of autoimmunity, cancer and pathogen infection (10–17).

Recently, a breadth of structural data has provided crucial insights into the mechanism of MHC-I chaperone function. Two crystal structures of TAPBPR-bound empty MHC-I molecules have illuminated aspects of the chaperoning and peptide editing functions (18)(19). TAPBPR has fused β-sandwich and immunoglobulin-like type 5 (IgV) folds forming a bilobed N-terminal domain with a large concave surface that cradles the underside of the α_1_α_2_ “platform”, beneath the MHC-I α_2_ domain. The peptide-binding groove is held in an open conformation at the F pocket, where the antigenic peptide C-terminus would normally reside, with smaller distortions extending to the A and B pockets. These groove perturbations promote the release of low affinity peptides. The C-terminal IgC domain of TAPBPR makes additional contacts to the α_3_/β_2_m domain junction, giving the appearance of a long scaffold supporting the different structural elements of the MHC-I. In one of the TAPBPR crystal structures (19), a protruding loop (a.a. G24-R36) of TAPBPR (**Fig. 1A**) was modeled in an α-helical conformation extending from the luminal tip of the N/IgV domain into the peptide-binding groove to occupy the F pocket (**Fig. 1B, C-middle**). The loop was hypothesized to “scoop” bound peptide out of the groove through the seductively simple mechanism of steric competition. A simultaneous publication from the same group of the tapasin-dependent PLC at low resolution seemed to support this mechanism, with the equivalent loop of tapasin partially modeled projecting into the F pocket (20) (**Fig. 1B, C-left**), while X-ray structures of MHC-I/dipeptide complexes together with a short hydrophobic peptide occupying the groove, derived from the tapasin loop, appear to show a stable, bound conformation (21). Recent studies have provided evidence that the TAPBPR G24-R36 loop influences peptide exchange on properly conformed MHC-I molecules, either by modulating peptide off-rates (22) or by acting as a leucine lever (23). In contrast, mutational insults to the “scoop” loop have only minor effects on tapasin-mediated intracellular processing of nascent MHC-I (21), while loop density is lacking in an independent crystal structure of the TAPBPR/MHC-I complex (18), likely due to disorder (**Fig. 1C-right**). Thus, while the TAPBPR G24-R36 loop seems to influence antigen selection, there remain outstanding questions of whether it may adopt a short helical conformation which enters the empty MHC-I groove, and how it affects peptide loading, relative to the shorter tapasin loop.

**Figure 1.**
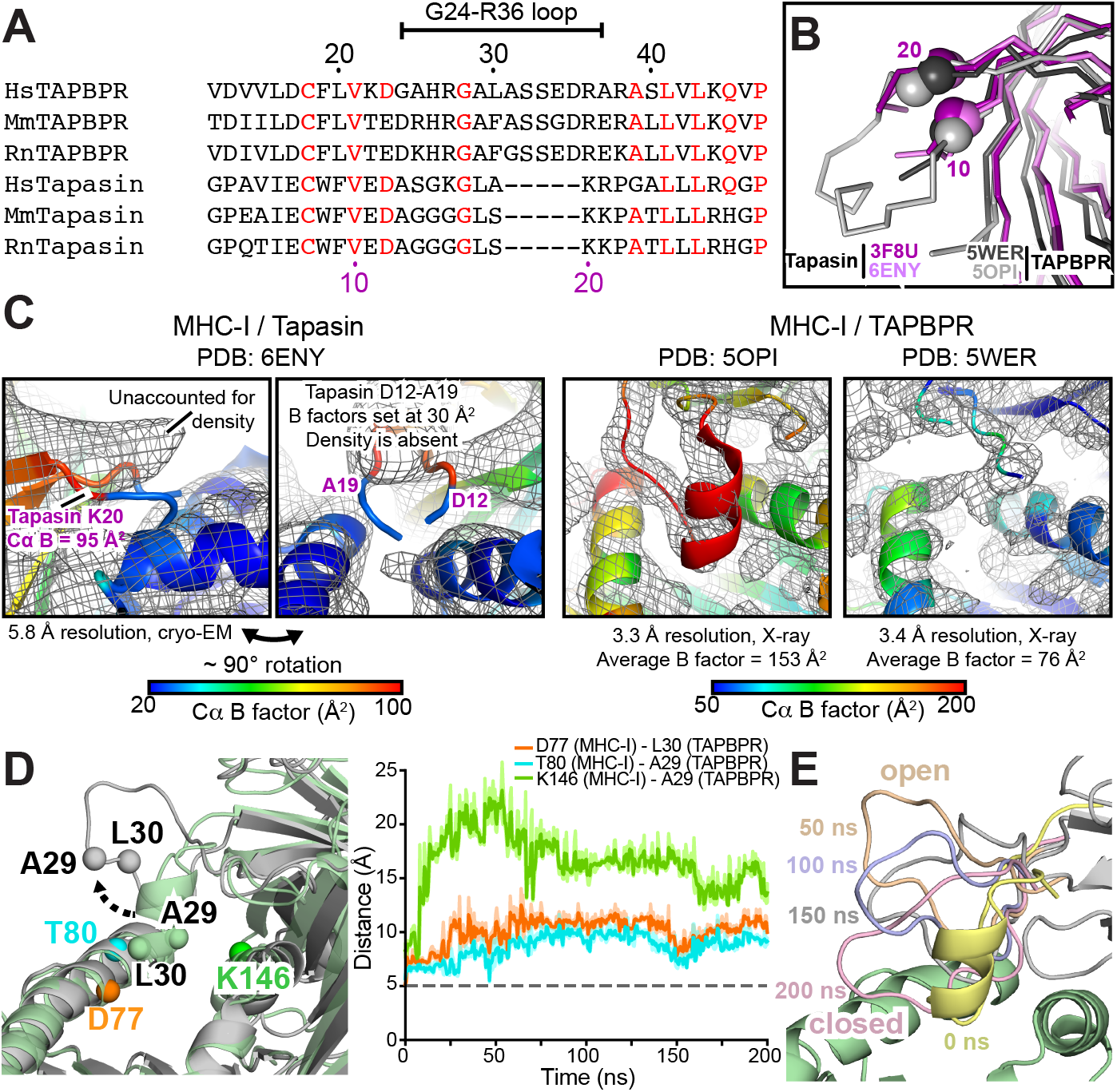
Conformational plasticity of the TAPBPR G24-R36 loop. (**A)** Sequence alignment between TAPBPR and tapasin from *Homo sapiens* (Hs, human), *Mus musculus* (Mm, mouse) and *Rattus norvegicus* (Rn, rat) highlighting differences in the TAPBPR G24-R36 loop region. Black and purple numbering reference TAPBPR and tapasin, respectively. Red residues are conserved. **(B)** Overlay of X-ray structures of tapasin (PDB IDs 3F8U and 6ENY) and TAPBPR (PDB IDs 5WER and 5OPI). **(C)** (Left) Cryo-EM density map (gray mesh, 5.0 σ) plotted on the cartoon of MHC-I in complex with tapasin (PDB ID 6ENY). (Right) 2Fo-Fc electron density maps (gray mesh, 1.0 σ) plotted on the cartoon of MHC-I in complex TAPBPR (PDB ID 5OPI and 5WER). The cartoons are colored based on the Cα B factors. (**D)** (Left) Before (green) and during (gray) snapshots of peptide-deficient HLA-A*02:01/hβ_2_m/TAPBPR complex from all-atom molecular dynamics simulations. The Cα atoms used for distance measurements are shown as spheres on the structure. (Right) Intermolecular Cα-Cα distances measured between HLA-A*02:01 groove (D77, T80, K146) and TAPBPR G24-R36 loop (A29, L30) residues over the course of the simulation. The dotted line represents Cα-Cα distances at the start of the simulation. **(E)** The range of conformations of the TAPBPR G24-R36 loop captured at different times during the MD simulation. The open (wheat, 50 ns) and closed (pink, 200 ns) states of the TAPBPR G24-R36 loop are oriented away from and towards the HLA-A*02:01 groove, respectively.

Here, using a combination of *in situ* functional assays, yeast display/MHC-I tetramer staining, and a range of complementary biophysical techniques including solution NMR, we address discrepancies in the MHC-I/TAPBPR X-ray structures and determine the role of the TAPBPR G24-R36 loop in the chaperoning and editing functions. Our findings show that the G24-R36 loop is not necessary for chaperoning activity, but directly participates in peptide editing on the MHC-I groove. Surprisingly, the TAPBPR G24-R36 loop does not interact directly with the floor of the MHC-I groove to compete with incoming peptides. Instead, the loop hovers above the MHC-I groove to promote loading by acting as a trap that occludes peptide dissociation. Together, our results underscore a role for TAPBPR as an auxiliary editor that operates in compartments of reduced peptide concentration.

## Results

### Unexpected conformational plasticity of the TAPBPR G24-R36 loop

A re-examination of the PLC cryo-EM structure(20) shows that the tapasin loop modeled as projecting in to the F pocket falls outside the electron density and has B factors fixed at 30.00 Å^2^, indicating it was excluded from refinement. Nearby density suggests that the loop instead rests above the peptide-binding groove (**Fig. 1C**). The TAPBPR G24-R36 loop was built in only one of the TAPBPR co-crystal structures where it was bound to the H-2D^b^ murine MHC-I (19), and its assigned B factors were exceedingly high (**Fig. 1C**; PDB ID 5OPI), with additional large B factor transitions between bonded amino acids across the entire model. In the second crystal structure of TAPBPR-bound H-2D^d^ (18), no clear density was observed for the 24-36 loop in the 4 different complexes of the asymmetric unit (**Fig. 1C**; PDB ID 5WER). The loop’s conformation and placement relative to the empty MHC-I groove are therefore uncertain in the original structural data.

To investigate the structural plasticity of the TAPBPR G24-R36 loop in the MHC-I/TAPBPR complex, we used complementary computational approaches. We chose the most common human MHC-I molecule, HLA-A*02:01, as our model system. In the first approach, the HLA-A*02:01 sequence was threaded onto the H2-D^d^/TAPBPR X-ray structure missing electron density for the G24-R36 loop (PDB ID 5WER) (18). The resulting HLA-A*02:01/TAPBPR complex was used as a template to explore whether an α-helical conformation could be sampled during loop modeling. We employed a variety of sampling algorithms, including cyclic coordinate descent (CCD), kinematic closure (KIC), and comparative modeling (CM), within the *Rosetta* modeling suite (24). The resulting lowest energy HLA-A*02:01/TAPBPR models revealed a wide range of TAPBPR G24-R36 loop conformations, each differing significantly from the α-helical conformation of the H-2D^b^/TAPBPR refined X-ray structure (**Fig. S1A-C**, **Fig. 1A-C**). In the second approach, the HLA-A*02:01 sequence was threaded onto the H2-D^b^/TAPBPR X-ray structure containing the G24-R36 “scoop loop” modeled with poorly resolved electron density (PDB ID 5OPI). The modeled HLA-A*02:01/TAPBPR complex was used as a starting point to determine whether the “scoop loop” conformation was stable in all-atom molecular dynamics (MD) simulations, performed in explicit solvent. Our MD trajectory revealed a loss of helical propensity of the short α-helix formed by TAPBPR residues G24-L30, as well as movement of the TAPBPR G24-R36 loop away from the HLA-A*02:01 groove within a 50 nsec timescale. This was highlighted by a marked increase in Cα-Cα distances measured between HLA-A*02:01 groove (D77, T80, K146) and TAPBPR G24-R36 loop (A29, L30) residues over the course of the simulation (**Fig. 1D, E**). During the MD simulations the TAPBPR G24-R36 loop sampled both an open state (pointing away from MHC-I groove) and a closed state that hovers above the MHC-I groove in an extended conformation (**Fig. 1D, E**). MD simulations and *Rosetta* modeling use different energy functions, and yet the two approaches independently sampled similar TAPBPR G24-R36 loop conformations (**Fig. S1A-C, Fig. 1D, E**). These results suggest that the loop is not constrained in an α-helical structure through interactions with the MHC-I groove. Instead, the TAPBPR G24-R36 loop may adopt an ensemble of non-helical conformations which do not interact with the MHC-I groove.

### TAPBPR chaperoning of nascent MHC-I molecules is defined by scaffolding domains

Tapasin was knocked out in human Expi293F cells, which caused diminished chaperone-dependent processing and trafficking of endogenous HLA-A*02:01 to the cell surface, as measured by flow cytometry (**Fig. S2 and S3A**). Surface expression of HLA-A*02:01 was rescued by transfection with a plasmid encoding tapasin (**Fig. 2A, S2 and S3A**). Overexpression of TAPBPR also partially rescued surface HLA-A*02:01 levels (**Fig. S3A**). Based on structural alignments (**Fig. 1B**), residues 11-19 of the tapasin loop were replaced with 22-35 of TAPBPR, and the protein remained functional, albeit with decreased activity which may reflect a minor structural perturbation (**Fig. 2A**). G15 of tapasin resides at the very tip of the loop, and shortening the loop by two residues either side (tapasin Δ2step) had no effect on tapasin activity (**Fig. 1A and 2A**). Shortening the loop further to remove L18 (tapasin Δ3step) caused a sudden drop in activity, although the protein still remained highly active. The cryo-EM density for the PLC (20) is consistent with tapasin L18 contacting the rim of the peptide-binding groove. Substitutions of tapasin L18 caused similar decreases in activity, whereas substitutions of G15 at the very tip of the loop had minimal effect. The data show that for folding and processing of HLA-A*02:01, a long loop on the chaperone to act as a scoop is not necessary, but binding contacts to the top of the MHC-I α-helices are important.

**Figure 2.**
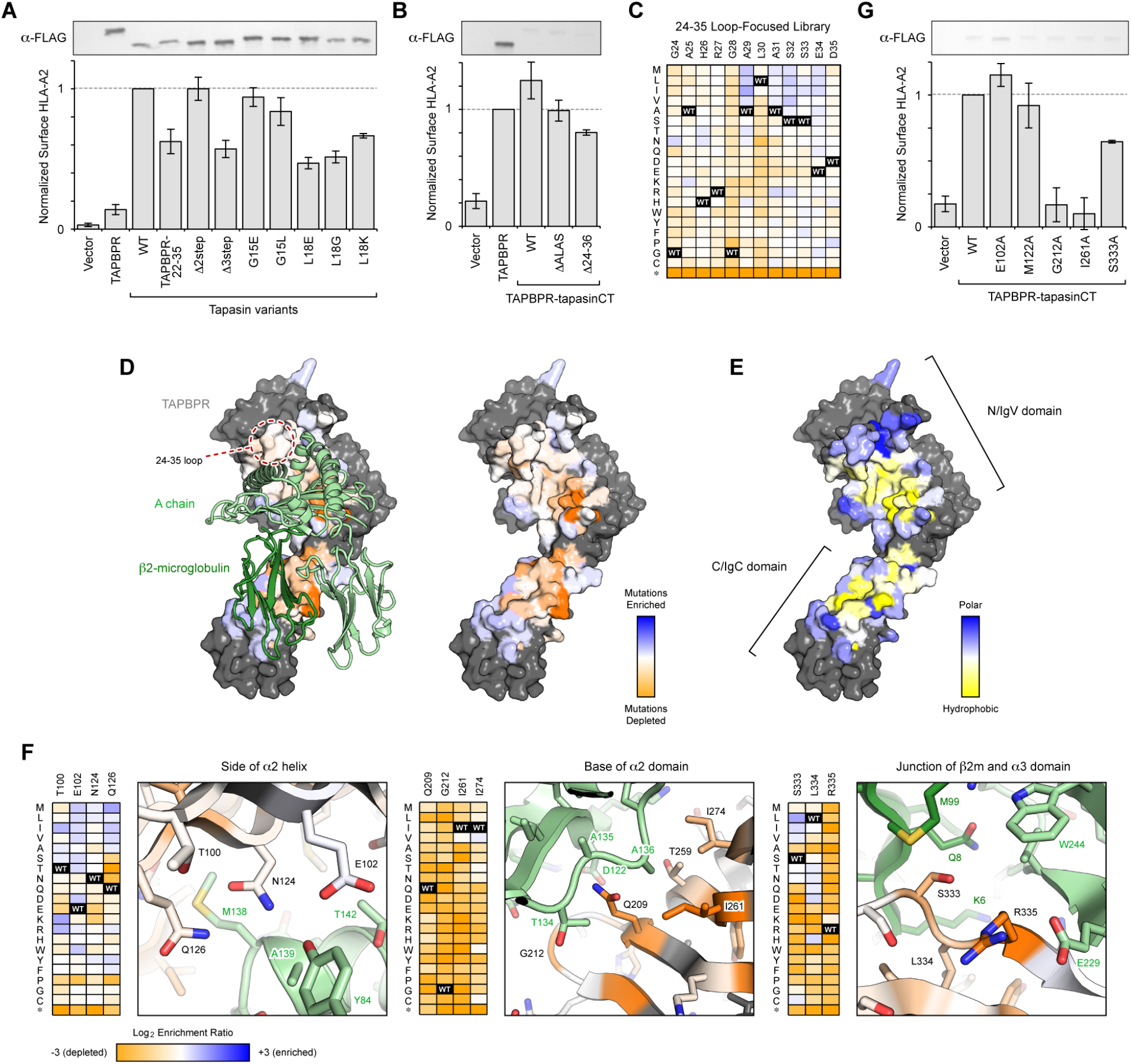
Important sequence features of TAPBPR for functional replacement of tapasin localize to scaffolding sites of MHC-I. **(A)** TAPBPR and variants of tapasin were transfected in tapasin-KO Expi293F cells, and relative surface HLA-A*02:01 expression as measured by flow cytometry is plotted. Data are mean ± SD, n = 4. Proteins were FLAG-tagged at their luminal N-termini, and immunoblots of whole lysate are shown in the upper inset. **(B)** A chimera of the TAPBPR luminal domain with the tapasin TM and cytosolic domains, called TAPBPR-CT, has increased activity for chaperoning endogenous HLA-A*02:01, despite reduced protein expression by immunoblot (*upper inset*). Variants of TAPBPR were tested for rescue of surface HLA-A*02:01 in the TAPBPR-CT background. Data are mean ± SD, n = 4. **(C)** A deep mutational scan of the TAPBPR 24-35 loop based on selection of tapasin-KO cells with rescued surface HLA-A*02:01. Log_2_ enrichment ratios for each mutation are plotted from ≤ −3 (depleted/deleterious, orange) to ≥ +3 (enriched, dark blue). TAPBPR residue position is on the horizontal axis, and amino acid substitutions are on the vertical axis (*, stop codon). **(D)** Conservation scores from an expanded deep mutational scan of the entire TAPBPR/MHC-I interface are mapped to a homology model of HLA-A*02:01-bound TAPBPR. Highly conserved residues on the TAPBPR surface for mediating the folding and surface trafficking of HLA-A*02:01 are colored orange, while neutral regions are pale white/blue. Residues excluded from the library and analysis are grey. MHC-I H chain and hβ_2_m are pale and dark green ribbons, respectively. **(E)** Log_2_ enrichment ratios are weighted by Kyte-Doolittle hydrophobicity and averaged for all substitutions at each residue position. The scores are mapped to the surface of TAPBPR, with residues having a preference for hydrophobics in yellow to polars in dark blue. **(F)** Close up views of three structural regions, colored as in D. Accompanying heatmaps plot Log_2_ enrichment ratios for each mutation from depleted/deleterious (orange) to neutral (white and pale colors) to enriched (dark blue). **(G)** Individual mutations of TAPBPR-CT were validated by targeted mutagenesis in the tapasin surrogate assay. Data are mean ± SD, n = 4. Total protein expression is shown in the upper anti-FLAG immunoblot.

We refer to the functional replacement of tapasin with TAPBPR in tapasin-knockout cells as a tapasin surrogate assay. Overexpression of TAPBPR partially rescued surface HLA-A*02:01 levels, although a subset of cells retained HLA-A*02:01 intracellularly at even greater levels (**Fig. 2A and S3A**). Substitution of the TAPBPR transmembrane and cytosolic segments with those of tapasin (called TAPBPR-CT) increased TAPBPR-mediated rescue of surface HLA-A*02:01, despite substantially reduced protein expression (**Fig. 2B**). We hypothesize that TAPBPR-CT may be recruited in to the PLC (**Fig. S2**), though this remains undemonstrated. TAPBPR-CT was used as the background in which mutations to the luminal domains of TAPBPR were tested. Deletion of the TAPBPR loop tip (TAPBPR ΔALAS, removing the last turn of the helix modeled for the loop in PDB 5OPI) caused a small decrease in HLA-A*02:01 processing, and deletion of the entire TAPBPR 24-36 loop (ΔG24-R36) caused a further decrease in activity. These results demonstrate that the loop is not essential for chaperoning function, although some effects can be observed for different length and sequence variants.

The luminal domains of TAPBPR can be highly expressed as soluble monomeric protein for biophysical characterization, and therefore TAPBPR became our focus for further investigation. Using the tapasin surrogate assay as the basis for fluorescence-based selection, the TAPBPR 24-35 loop was deep mutationally scanned to assess the relative activities of all 240 single amino acid substitutions. Apart from heavy depletion of nonsense mutations, the mutational landscape across the 24-35 loop is relatively featureless (**Fig. 2C and S3C**), consistent with a nonessential role in HLA-A*02:01 processing.

To compare with other regions of TAPBPR that may be important, a larger library was constructed encompassing all single amino acid substitutions across 75 TAPBPR residues at the crystallographic interface with MHC-I (18). As controls, saturation mutagenesis was also applied to 10 surface residues on the opposite side and to 7 buried residues expected to be important for TAPBPR folding. Following selection of the TAPBPR library for high levels of surface HLA-A*02:01 expression (**Fig. S3D-S3F**), mutations to buried residues in the TAPBPR core were found to be depleted except for hydrophobic substitutions, whereas surface residues distal from the MHC-I interface were mutationally permissive (**Fig. S3G**). There were two striking regions of TAPBPR within the MHC-I interface that were tightly conserved for activity: *(i)* the concave face of the N/IgV domain that rests below the MHC-I α_2_ domain, and *(ii)* the edge of the C/IgC domain that bridges the β_2_m and α_3_ domains (Fig. 2D). Weighting the enrichment ratios by hydrophobicity suggests that most interactions in these regions have a preference for mutations to hydrophobic residues (**Fig. 2E**). By comparison, TAPBPR residues proximal to the upper rim of the peptide-binding groove showed higher tolerance towards mutation (**Fig. 2F**). Deep mutational scans achieve scale at the cost of data noise, and targeted alanine substitutions were tested to confirm that TAPBPR residues contacting the α_2_ underside (G212A and I261A) and α_3_/β_2_m junction (S333A) were critical for mediating rescue of surface HLA-A*02:01 in tapasin-knockout cells, whereas residues contacting the upper rim of the peptide groove (E102A and M122A) were not (**Fig. 2G**). Overall, critical sequence features of TAPBPR map to sites that scaffold the correct folded architecture of the heavy chain with β_2_m, consistent with our prior conclusions that TAPBPR chaperones nascent, improperly folded MHC-I substrates within the cell(5). Notably, the G24-R36 loop is not critical for this activity.

### TAPBPR recognition of properly conformed pMHC-I is affected by the G24-R36 loop

Trafficking of HLA-A*02:01 to the plasma membrane will depend on chaperoning activity of TAPBPR, but not necessarily its editing function. To directly assess how mutations impact TAPBPR binding to folded pMHC-I molecules, as would occur during editing of the bound peptide, the extracellular domain of TAPBPR was displayed on the surface of yeast and binding to fluorescent pMHC-I tetramers was detected by flow cytometry (**Fig. S4A**). Bound MHC-I was competed off by the addition of peptides in a dose-dependent manner that correlated with peptide affinities (**Fig. S5D and S5E**). The TAPBPR 24-35 loop was mutagenized, and the yeast-displayed library was screened for tight binding to two MHC-I alleles: mouse H2-D^d^ and human HLA-A*02:01 (**Fig. S4B**). No binding was observed for tetramers of a third allele, human HLA-A*01:01, consistent with TAPBPR having restricted recognition of folded pMHC-I allotypes(25). Addition of peptides increased stringency of the selections, but did not change the overall form of the mutational landscapes (**Fig. S4C-G**). At nearly all positions of the 24-35 loop there was enrichment of large, hydrophobic and aromatic residues for increased binding to both H2-D^d^ and HLA-A*02:01 (**Fig. 3A**). Design of protein-protein interfaces has shown that large hydrophobic residues can promote binding through non-specific non-polar interactions (26)(27). Instead, our attention was drawn to allele-specific differences that are unlikely to be due to generic mutational effects. For binding to H2-D^d^, positions 28 and 30 near the center of the TAPBPR 24-35 loop have strong preferences for aliphatic or aromatic side chains, whereas all non-basic substitutions of R27 are enriched (excluding cysteine, which is deleterious at all positions). The data are therefore suggestive that the tip of the loop is near an hydrophobic patch on H2-D^d^ with a nearby electropositive charge; surfaces meeting these criteria can be found near the middle of the α_1_ helix (MHC-I residues V76 and R79) or at the end of the α_2-1_ helix (K146, A149, and A150). However, permissiveness in the landscape also suggests contacts between the 24-35 loop and H2-D^d^ are sufficiently loose to tolerate mutations. Amino acid preferences within the 24-35 loop were weaker in the selections for binding HLA-A*02:01, although position 30 again prefers hydrophobic side chains. L30 has also been found to be important for binding to HLA-A*68:02(23). TAPBPR D35 is tightly conserved as an acidic residue for high H2-D^d^ binding due to electrostatic complementarity with basic residues (H2-D^d^ R79 and R83) on the upper rim of the peptide-binding groove. HLA-A*02:01 has a glycine at position 79, and nearly all non-acidic substitutions of TAPBPR E34 and D35 enhance HLA-A*02:01 binding, consistent with Rosetta models of the 24-35 loop contacting the upper surfaces of the MHC-I a-helices (**Fig. 3B**).

**Figure 3.**
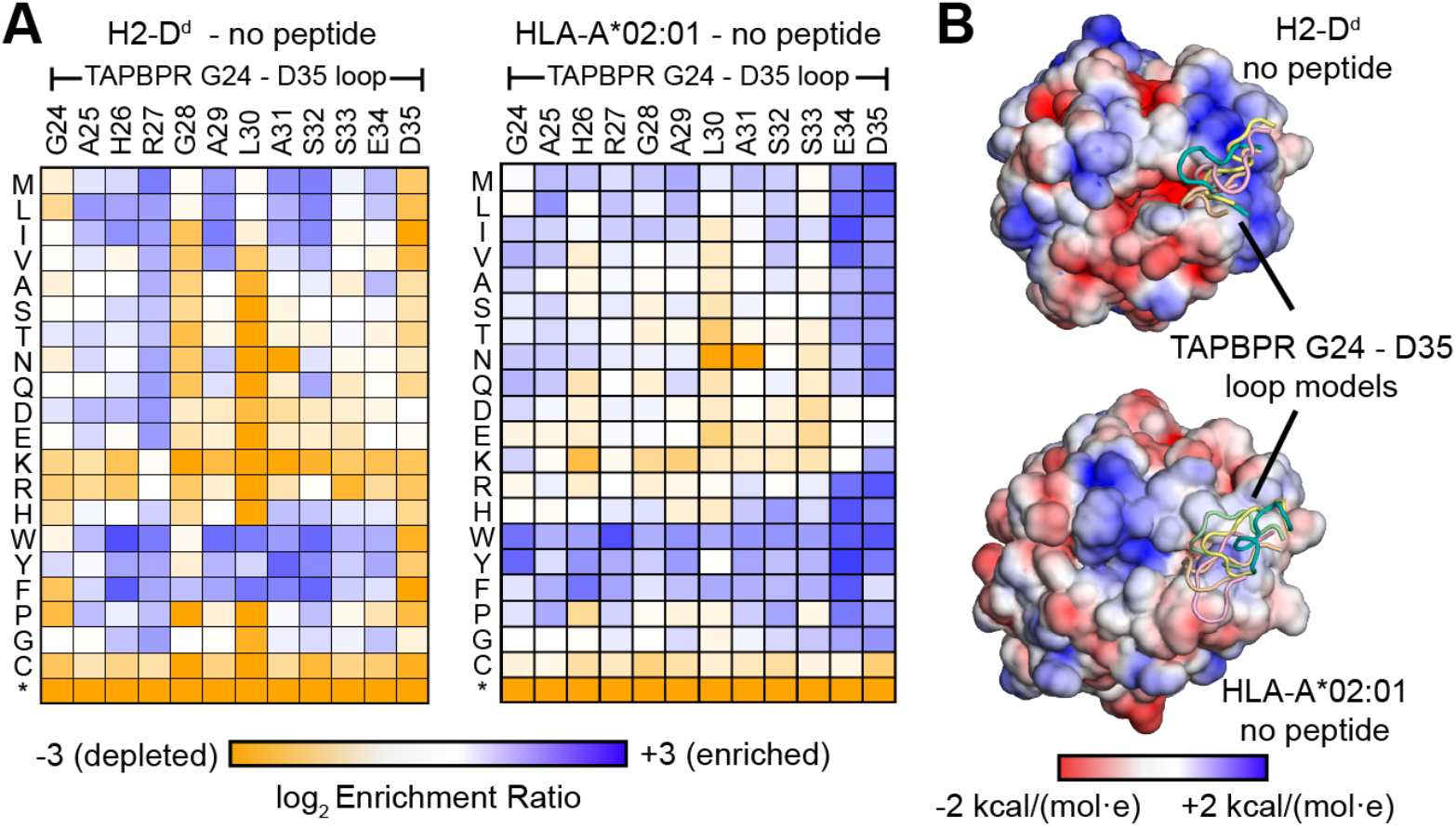
TAPBPR 24-35 loop sequence interaction landscape is pMHC-I allele-dependent. **(A)** A TAPBPR library of substitution mutants in the 24-35 loop was displayed on the yeast surface and sorted for high binding signal to fluorescent pMHC-I tetramers made using refolded P18-I10/H2-D^d^ and TAX9/HLA-A*02:01 as indicated. Positive selection refers to tetramer staining under no excess of competing peptide, aiming to enrich for sequence variants of improved binding to the MHC-I tetramers. In the mutational landscapes, Log_2_ enrichment ratios for each mutation are plotted from ≤ −3 (depleted/deleterious, orange) to ≥ +3 (enriched, dark blue). TAPBPR 24-35 loop residue position is on the horizontal axis, and amino acid substitutions are on the vertical axis (*, stop codon). **(B)** Solvent-accessible surfaces of MHC-I molecules colored by electrostatic potential (negative in red, to positive in blue) as calculated in CHARMM-GUI PBEQ-Solver for H2-D^d^ (PDB 3ECB) and HLA-A*02:01 (PDB 1DUZ). The five lowest energy RosettaCM models of the TAPBPR 24-35 loop are shown as references.

Selected mutations that increase or decrease MHC-I binding, predicted from the deep mutational scans, were validated by targeted mutagenesis (**Fig. S5**). Furthermore, the binding of yeast-displayed TAPBPR DALAS to tetramers of H2-D^d^ or HLA-A*02:01 was only slightly reduced compared to wild type (WT). TAPBPR ΔALAS and targeted point mutants exhibited competitive interactions with peptide (**Fig. S5D and S5E**), indicating that differences in how tightly variants of the G24-R36 loop recognize MHC-I has no major bearing on peptide-mediated chaperone displacement. Rather, other sites of contact between TAPBPR and MHC-I must act as the sensors for high affinity peptide binding within the groove. We further note that peptides of different sequence were slightly better or worse at displacing some TAPBPR mutants, indicating mutations in the G24-R36 loop have subtle, peptide sequence-specific effects.

### The TAPBPR loop does not enter the empty MHC-I groove

TAPBPR mutants ΔALAS and ΔG24-R36 were recombinantly expressed in S2 *Drosophila* cells, purified and characterized *in vitro*. Circular dichroism (CD) spectroscopy shows that TAPBPR G24-R36 loop mutants retain the expected immunoglobulin (Ig)–like fold, as highlighted by the 2-sheet characteristic of a negative band between 215 and 219 nm (**Fig. 4A, top**). By differential scanning fluorimetry (DSF), TAPBPR G24-R36 loop mutants exhibit thermal stability comparable to WT with melting temperature (T_m_) in the 48-49 °C range (**Fig. 4A, bottom**). Finally, both TAPBPR^WT^ and TAPBPR^ΔG24-R36^ proteins readily form high-affinity complexes with empty H2-D^d^ or HLA-A*02:01 molecules, prepared upon refolding with the photosensitive peptides photoP18-I10 and photoFluM1, respectively, upon UV-mediated peptide release (**Fig. S6A-B)**.

**Figure 4.**
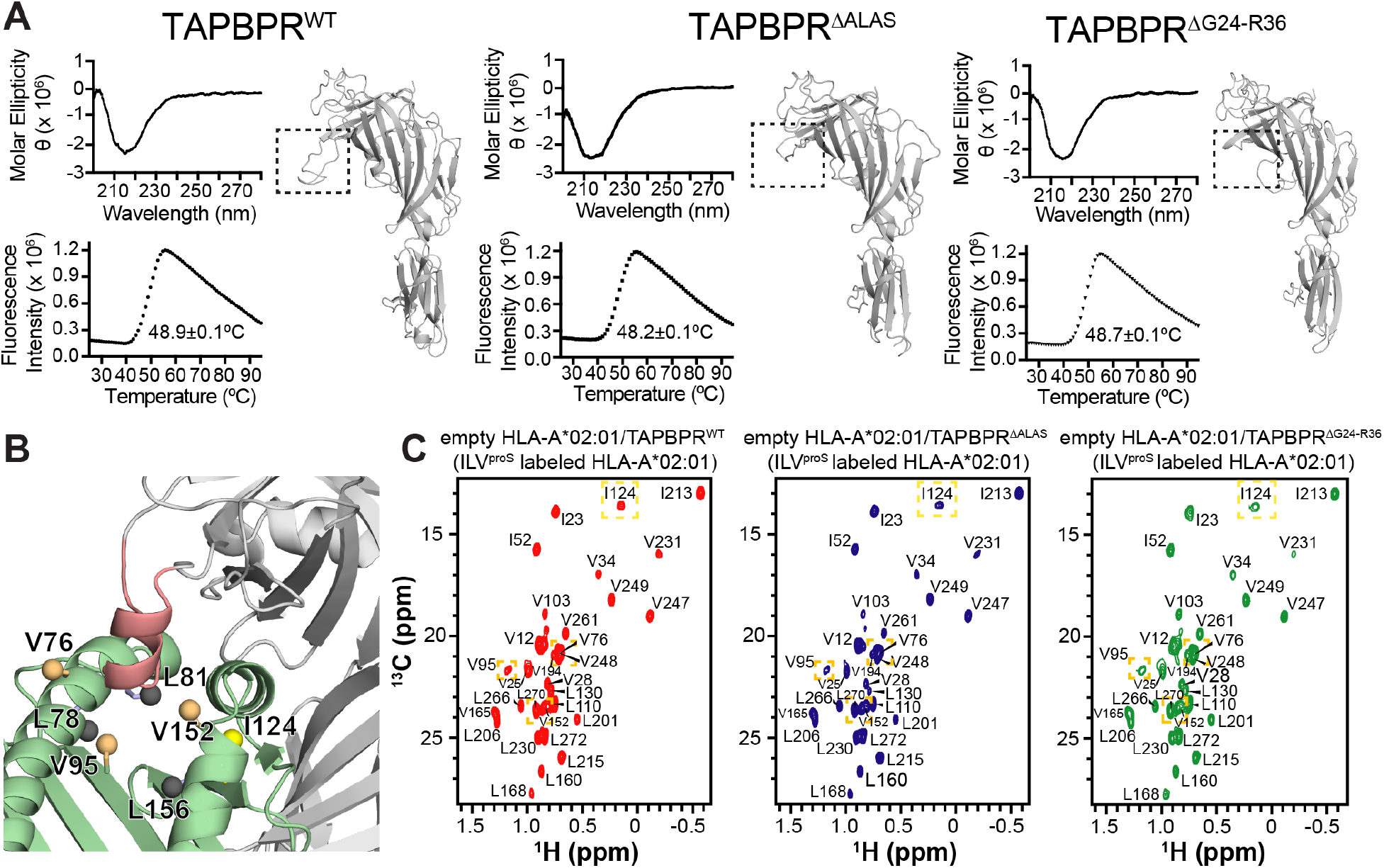
The TAPBPR G24-R36 loop does not enter the HLA-A*02:01 groove. **(A)** Comparison between wild-type (WT) and mutant TAPBPR constructs used in this study. ΔALAS = deletion of residues A29-S32 and ΔG24-R36 = deletion of residues G24-R36. (Top) Far-UV CD and (Bottom) DSF spectra of each TAPBPR construct. The average from three technical replicates is shown. An inset in the DSF spectra notes measured thermal melt (Tm) values. The corresponding *RosettaCM* model of each TAPBPR construct is shown. The dotted box highlights differences in the G24-R36 loop region. **(B)** View of the peptide-deficient HLA-A*02:01/TAPBPR model (template PDB ID 5OPI) showing ILV^proS^ methyl probes on HLA-A*02:01 (as spheres) within 10 Å of the TAPBPR G24-R36 loop (salmon). Methyl resonances of residues L78, L81 and L156 (shown in black) are missing in 2D ^1^H-^13^C methyl HMQC spectra of peptide-deficient HLA-A*02:01/TAPBPR complex due to conformational exchange induced line broadening. **(C)** 2D ^1^H-^13^C methyl HMQC spectra of 80 μM peptide-deficient HLA-A*02:01 (ILV^proS^ labeled)/hβ_2_m in complex with TAPBPR^WT^ (red), TAPBPR^ΔALAS^ (blue) or TAPBPR^ΔG24-R36^ (green) recorded at 25°C at a ^1^H field of 800 MHz. Dotted boxes highlight methyl probes (shown in panel B) that are modeled to be near the TAPBPR G24-R36 loop.

To investigate specific loop interactions with MHC-I molecules, we used solution NMR and characterized HLA-A*02:01/TAPBPR complexes prepared using either wild-type or mutant TAPBPR(5, 28). We isolated 1:1 peptide-deficient HLA-A*02:01/TAPBPR complexes where the HLA-A*02:01 heavy chain was selectively ILV^proS 13^C/^1^H methyl labeled on a ^12^C/^2^H background, whereas hβ_2_m and TAPBPR were at natural isotopic abundance. These experiments allowed us to examine how the presence of the TAPBPR G24-R36 loop influences the chemical environment of the 37 ILV^proS^ methyl probes on HLA-A*02:01, several of which are in the groove and within 10Å from the TAPBPR G24-R36 loop in the proposed “scoop loop” conformation (**Fig. 4B**). Analysis could not be performed on a subset of the probes (L78 δ2, L81 δ2 and L156 δ2) due to NMR line broadening resulting from conformational exchange in the intermediate (μsec-msec) timescale (**Fig. 4B, gray spheres**). Chemical shift analysis of the remaining probes did not identify any noticeable changes in 2D ^1^H-^13^C methyl HMQC spectra of peptide-deficient HLA-A*02:01/TAPBPR complexes prepared with TAPBPR^WT^, TAPBPR^ΔALAS^, or TAPBPR^ΔG24-R36^ (**Fig. 4C**). These results were corroborated by similar observations using independently prepared peptide-deficient MHC-I/TAPBPR complexes employing a more general methyl labeling scheme (AILV) on HLA-A*02:01, where the number of total methyl probes increased to 94, 20 of which were within 10Å from the TAPBPR G24-R36 loop (**Fig. S7A**). We examined potential chemical shift changes in HLA-A*02:01/TAPBPR complexes bound to unlabeled TAX9 peptide, where the resonances of most HLA-A*02:01 methyl probes that were broadened in the peptide-deficient complexes became readily visible (**Fig. S7B)**. Chemical shift analysis of the AILV methyl probes in wild-type and mutant TAX9/HLA-A*02:01/TAPBPR complexes did not reveal any measurable changes in 2D ^1^H-^13^C HMQC spectra (**Fig. S7C**). Together, these findings support the lack of a direct, stable interaction between the TAPBPR G24-R36 loop and the HLA-A*02:01 groove either in the empty or the peptide-bound states.

### The TAPBPR loop promotes peptide loading on empty MHC-I molecules

We have previously characterized the full thermodynamic cycle of TAPBPR-mediated MHC-I peptide exchange (28). The cycle is defined by dissociation constants (K_D_) for four reversible steps: association of TAPBPR with MHC-I in the absence of peptide (K_D1_), association of peptide with MHC-I in the absence of TAPBPR (K_D2_), association of pMHC-I with TAPBPR (K_D3_), and association of peptide with the MHC-I/TAPBPR complex (K_D4_) (**Fig. 5A**). We measured apparent K_D_ values for each of the specific steps using suitable isothermal titration calorimetry (ITC) experiments. ITC was performed on a range of known HLA-A*02:01 peptides (TAX8 - LFGYPVYV, TAX9 - LLFGYPVYV and KLL15 - KLLEIPDPDKNWATL) to uncover any differences in the TAPBPR G24-R36 loop effect in relation to peptide length (29, 30). Optimization of ITC experimental conditions (*Materials and Methods*) allowed us to focus on different steps of the thermodynamic cycle (**Fig. 5B** and **Fig. S8A-C**). First, direct measurement of peptide to the empty MHC-I groove (K_D2_) can be limited by the inherent instability of peptide-deficient MHC-I molecules. Thus, we utilized TAPBPR as a stabilizer of peptide-deficient HLA-A*02:01. Purified 1:1 HLA-A*02:01/TAPBPR complex in the calorimeter cell was titrated by injecting peptide. The apparent K_D2_ obtained from the experiment includes the release of TAPBPR (**Fig. 5B-left** and **Fig. S8A)**. The K_D2_ values, which range from 40 to 400 nM across our peptide set, did not differ significantly between HLA-A*02:01/TAPBPR complexes prepared using TAPBPR^WT^ or TAPBPR^ΔG24-R36^, suggesting that our measurements were dominated by peptide binding to empty, unchaperoned MHC-I (**Fig. 5C** and **Table 1**). By titration of TAPBPR with pMHC-I in the presence of excess peptide in both the cell and syringe, the K_D3_ for the formation of ternary complex was obtained (**Fig. 5B-middle** and **Fig. S8A)**. Measured K_D3_ values, which were in the range of 2 μM for the different peptides in our set, were similar between pMHC-I/TAPBPR complexes formed using TAPBPR^WT^ or TAPBPR^ΔG24-R36^ (**Fig. 5C** and **Fig. S8A-C** and **Table 1**), suggesting that the loop does not affect recognition of peptide-loaded MHC-I. Finally, titration of the HLA-A*02:01/TAPBPR complex by injecting the peptide in the presence of an excess TAPBPR in both the cell and syringe (to minimize dissociation of TAPBPR from the pMHC-I/TAPBPR complex) allowed measurement of an apparent value for K_D4_, binding of peptide to an empty MHC-I/TAPBPR complex (**Fig. 5B-right** and **Fig. S8B)**. Here, we observed binding to two sequential sites (**Fig. 5B-right**), due to a combination of processes that are described by the steps indicated with K_D2_ and K_D4_ (**Fig. 5A**) under our working protein concentrations. However, given that the K_D2_ process does not involve TAPBPR, results obtained using different TAPBPR form can be used to probe effects of the loop on K_D4_.

**Table 1.**
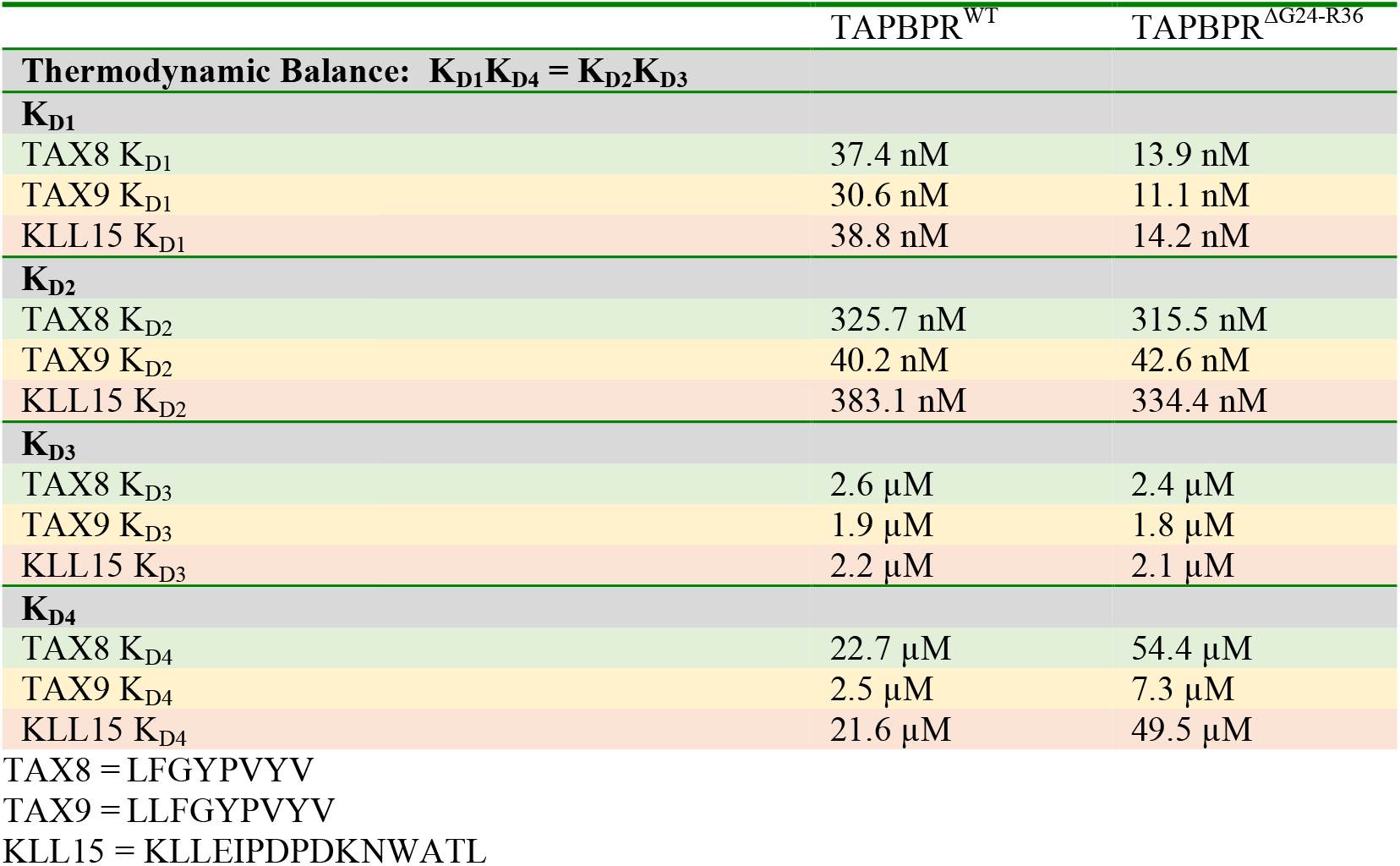
ITC derived apparent dissociation constants for each step of the TAPBPR-mediated peptide exchange cycle

**Figure 5.**
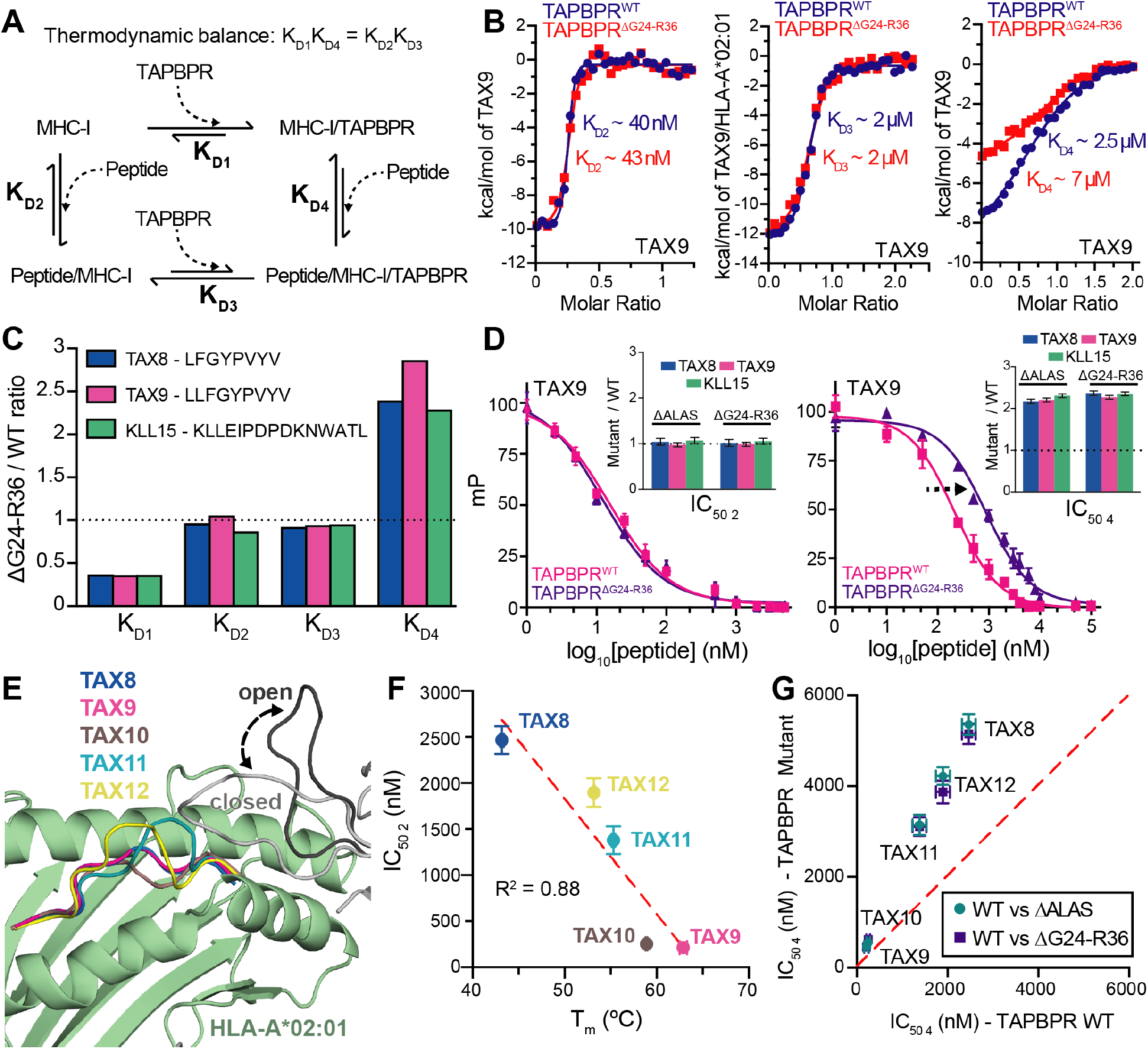
The TAPBPR G24-R36 loop promotes peptide binding on empty MHC-I. **(A)** Schematic of the TAPBPR-mediated MHC-I peptide exchange cycle. The dissociation constant (K_D_) of each step is noted. **(B)** ITC performed at different stages of the peptide exchange cycle (K_D2_, K_D3_, K_D4_) for TAX9/HLA-A*02:01/hβ_2_m with TAPBPR^WT^ or TAPBPR^ΔG24-R36^ (*Materials and Methods*). K_D1_ was not measured directly, but inferred from thermodynamic balance along the cycle shown in (A). **(C)** Apparent K_D_ ratios determined by ITC for TAPBPR^ΔG24-R36^ / TAPBPR^WT^ for TAX8, TAX9 and KLL15 peptides. The dotted line represents no effect. (**D)** FP performed in the absence (left) or presence (right) of 20-fold molar excess [TAPBPR]. Millipolarization (mP) values are plotted as a function of the log10 peptide concentration for TAX8, TAX9 and KLL15 competitor peptides. Error bars were obtained from three technical replicates. The insets show a comparison of the ratio of FP determined IC_50_ values for TAPBPR^ΔG24-R36^ or TAPBPR^ΔALAS^ versus TAPBPR^WT^. The dotted line represents no effect. **(E)** X-ray structures (TAX8, TAX9) and *RosettaCM* models (TAX10, TAX11, TAX12) show peptide bulging within the HLA-A*02:01 groove. The open and closed conformation of the TAPBPR G24-R36 loop from MD simulations are shown in gray and black, respectively. **(F)** Comparison of FP determined IC_50_ values under non saturating TAPBPR concentrations for TAX length variants versus pMHC-I thermal stability. R^2^ values of the linear regression (red line) are shown. **(G)** Comparison of FP determined IC_50_ values under 20-fold molar excess of TAPBPR for TAX length variants for TAPBPR^WT^ versus TAPBPR^ΔALAS^ or TAPBPR^ΔG24-R36^. The dotted red line is a reference for a 1:1 correlation (no effect).

Notably, apparent K_D4_ values were increased by a factor of two to three-fold between pMHC-I/TAPBPR complexes prepared using TAPBPR^ΔG24-R36^ versus TAPBPR^WT^ (**Fig. 5A-C** and **Fig. S8A-C** and **Table 1**); i.e. the G24-R36 loop increases the affinity of empty, chaperoned MHC-I for incoming peptides. While instability of peptide-deficient HLA-A*02:01 molecules in the absence of TAPBPR did not permit direct measurement of K_D1_, binding of TAPBPR to empty MHC-I, this value was inferred from thermodynamic balance along the exchange cycle (**Fig. 5A** and **Table 1**). K_D1_ was decreased by a factor of two to three-fold for TAPBPR^ΔG24-R36^ versus TAPBPR^WT^ (**Table 1**), suggesting that the presence of the loop reduces the affinity of TAPBPR for empty MHC-I. This result is inconsistent with models of the TAPBPR 24-36 loop acting as a stabilizer of empty MHC-I molecules and a steric competitor of incoming peptides. In agreement with our NMR results, the ITC data suggest that the TAPBPR G24-R36 loop does not form a strong, stabilizing interaction with the empty MHC-I groove but instead functions to promote peptide loading to the empty MHC-I/TAPBPR complex.

### The TAPBPR loop length is important for *in vitro* peptide loading activity

To determine whether the full length of the TAPBPR G24-R36 loop is required for normal function, we applied a similar design approach from our ITC studies to fluorescence polarization (FP) experiments. We have previously shown that the association of peptides with the MHC-I groove can be monitored with FP competition assays where purified peptide-deficient MHC-I/TAPBPR complexes are incubated with fluorescently labeled TAMRA-peptide and a range of concentrations of unlabeled competitor peptide (28). Analogous to our ITC experiments, optimization of FP assay conditions (*Materials and Methods*) allowed us to independently probe each step of the thermodynamic cycle (**Fig. S9A-C**). First, dilution of peptide-deficient HLA-A*02:01/TAPBPR complexes in the absence of excess TAPBPR promotes dissociation of HLA-A*02:01 from TAPBPR upon titration of peptide, which allows for measurement of peptide IC_50 2_ values. FP was performed with TAX8, TAX9, and KLL15 serving as the competitor peptides. In agreement with ITC, IC_50_ 2 values determined by FP did not differ between HLA-A*02:01/TAPBPR complexes prepared using TAPBPR^WT^, TAPBPR^ΔALAS^, or TAPBPR^ΔG24-R36^ (**Fig. 5D, left** and **Fig. S9B**). Second, titration of TAPBPR into a sample containing fluorescently labeled TAMRA-TAX9 in complex with HLA-A*02:01 allows measurement of K_D3_ in the peptide exchange cycle (**Fig. 5A**). In agreement with our ITC experiments, we find that measured K_D3_ values from FP, which are in a similar range to ITC of 2 μM, did not differ between pMHC-I/TAPBPR complexes formed using TAPBPR^WT^, TAPBPR^ΔALAS^, or TAPBPR^ΔG24-R36^ (**Fig. S9A**). Finally, dilution of peptide-deficient HLA-A*02:01/TAPBPR complexes under in the presence of excess TAPBPR to minimize dissociation of TAPBPR from the pMHC-I/TAPBPR complex allows for measurement of IC_50_ 4. In agreement with our ITC data, we find that measured IC_50 4_ values are increased by a factor of two to three-fold between pMHC-I/TAPBPR complexes formed using ΔG24-R36 versus wild-type TAPBPR (**Fig. 5D, right** and **Fig. S9C**). In summary, these results provide corroborating evidence that the TAPBPR G24-R36 loop influences formation of the pMHC-I/TAPBPR intermediate complex. In addition, because we find that TAPBPR^ΔALAS^ with a short loop deletion has the same behavior as TAPBPR^ΔG24-R36^, our results suggest that the loop length is an important factor for peptide exchange.

### The TAPBPR loop does not explicitly control the length of bound peptides

We applied our FP experiments to determine whether the length of the MHC-I cargo influences G24-R36 loop mediated peptide selection. Since it is established that peptides of length longer than ten bulge out of the MHC-I groove (30, 31), we hypothesized that steric clashes between protruding peptide residues with the G24-R36 loop, hovering above the groove, could influence TAPBPR-mediated peptide exchange. To test our hypothesis, we prepared a set of peptides derived from the HTLV-1 TAX epitope ranging from 8 to 12 amino acids in length, each containing the same residue types in the two anchor positions. Comparison of X-ray structures available for TAX8 and TAX9 with *RosettaCM* models of TAX10 (LLFGGYPVYV), TAX11 (LLFGGGYPVYV), and TAX12 (LLFGGGGYPVYV) revealed the expected trend with longer peptides bulging from the HLA-A*02:01 groove **(Fig. 5E)**. Using DSF experiments we observed that optimal peptide length (9-10 residues (32)) correlates with highest thermal stability of the pMHC-I complex, as shown by the low stability of TAX8 and TAX12 (T_m_ = 43.2 and 53.2°C) and higher stability for TAX9 and TAX10 (T_m_ = 62.9 and 58.9°C) (**Fig. S10A**). Next, we performed FP competition experiments under conditions of low TAPBPR binding for each of the TAX length variants, to quantify peptide binding to empty MHC-I molecules upon release from TAPBPR. We observed a similar trend for FP determined IC_50 2_ values, where TAX9 and TAX10 peptides exhibited stronger association with HLA-A*02:01 (**Fig. S10B**). In addition, as observed in our ITC experiments, measured IC_50 2_ values were similar between HLA-A*02:01/TAPBPR complexes prepared using TAPBPR^WT^, TAPBPR^ΔALAS^, or TAPBPR^ΔG24-R36^ (**Fig. S10B**). We observed an excellent correlation between FP determined IC_50 4_ values and T_m_ values of pMHC-I complexes **(Fig. 5F)**. In addition, we measured IC_50 4_ values for the TAX peptide length variants and found an increase by a factor of 2-3 between pMHC-I/TAPBPR complexes prepared using TAPBPR^ΔALAS^ or TAPBPR^ΔG24-R36^ versus TAPBPR^WT^, consistently across all peptides (**Fig. 5G** and **Fig. S10C**). It is worth noting that the 15mer KLL15 peptide exhibits a similar G24-R36 loop effect compared to our TAX length variant dataset (**Fig. 5D, G**). Together, these data suggest that the TAPBPR G24-R36 loop promotes loading of all high-affinity peptides, irrespective of their length.

## Discussion

Recently, a breadth of structural data has provided invaluable insights into the mechanism of MHC-I chaperone function. These studies have revealed that chaperone mediated peptide exchange/editing is achieved through *(i)* stabilization of the peptide-deficient MHC-I in a peptide-receptive conformation, *(ii)* ejection of suboptimal peptides by inducing an ~3 Å widening of the MHC-I groove, and *(iii)* regulation of a dynamic switch located in the MHC-I groove (20, 28, 33, 34). Despite these important findings, the mechanistic details of how chaperones influence the repertoire of MHC-I presented antigens remains incompletely characterized, while significant differences between the two published X-ray structures challenge the proposed role of the TAPBPR G24-R36 loop as a direct peptide competitor.

Here we resolve key discrepancies between recent MHC-I/TAPBPR X-ray structures by examining the conformational landscape of the TAPBPR G24-R36 loop and directly probing for an interaction between the TAPBPR G24-R36 loop and the MHC-I groove using solution NMR. Our data suggest that the TAPBPR G24-R36 loop is intrinsically disordered, sampling a wide range of conformations adjacent to the MHC-I groove, but does not form stable contacts with the floor of the empty MHC-I groove **(Fig. 1D, E and Fig. 4C)**. These observations are consistent with the H2-D^d^/TAPBPR structure in which the TAPBPR G24-R36 loop is missing due to poorly defined electron density (likely as the result of conformational variation), rather than the H2-D^b^/TAPBPR structure with an ordered loop. Our data and interpretations are also consistent with electron density for the equivalent loop of tapasin in the PLC cryo-EM structure, yet for reasons that are not clear, the authors chose to model the loop outside of the density as entering the empty MHC-I groove. One limitation of our study is the lack of an HLA-A*02:01/TAPBPR X-ray structure, thus our study is restricted to interpreting models built from available X-ray structures as templates **(Fig. 1 and Fig. S1)**. In addition, while our solution NMR experiments provide clear evidence for a lack of a direct interaction between the TAPBPR G24-R36 loop and the MHC-I groove (**Fig. 4C**), these data were acquired on the human HLA-A*02:01 heavy chain and it remains unclear how differences in amino acid polymorphisms within the mouse H2 groove influence behavior of the TAPBPR G24-R36 loop (5, 8). While we did not find differences in the formation of empty H2-D^d^/TAPBPR or HLA-A*02:01/TAPBPR complexes *in vitro* (**Fig. S6**), our yeast display experiments suggest there are differences in behavior of the TAPBPR G24-R36 loop as the result of allele specific chemistry of the pMHC-I surface **(Fig. 3)**, in agreement with recent binding results using a range of human HLA alleles(35).

We sought to determine the role of the G24-R36 loop on TAPBPR’s recently identified dual functions of chaperoning and peptide editing. Deep mutagenesis based on the ability of TAPBPR to functionally replace tapasin for HLA-A*02:01 processing within the cell suggests that the G24-R36 loop does not substantially participate in the chaperone function. However, deep mutagenesis of yeast-displayed TAPBPR indicates the G24-R36 loop may have a specific role in interacting with folded pMHC-I, as occurs in peptide editing. To examine the role of the loop during editing, we carefully designed novel ITC and FP assays that allowed us to uniquely probe each step of the thermodynamic cycle describing the peptide exchange process **(Fig. 5A, S7 and S8)**. According to the “scoop loop” model, removal of the TAPBPR G24-R36 loop would result in a lower K_D4_ (tighter binding of peptides to the MHC-I groove). Instead, we find that removal of the TAPBPR G24-R36 loop results in a higher K_D4_ **(Fig. 5C, G)**. Together, these observations, in conjunction with our loop modeling and solution NMR studies, lead us to propose that the TAPBPR G24-R36 loop functions as a peptide trap (or lid), rather than a “scoop loop”.

We further examined whether the TAPBPR G24-R36 loop may have a function in selecting peptides with a specific length. Our data shows that the presence of the TAPBPR G24-R36 loop exerts a broad effect across the repertoire, independent on peptide length, by promoting binding of peptides according to their global affinity for the empty MHC-I groove **(Fig. S8 and S9)**, although some subtle effects were observed between TAPBPR loop mutants and specific peptide sequences by yeast display (**Fig. S5D**). In other words, while TAPBPR lowers the peptide affinity requirements across the sampled peptidome, it appears that the specificity of peptide binding is determined by interactions with the MHC-I groove, not the TAPBPR loop. It is conceivable that allele-specific deformations of the MHC-I groove, induced by TAPBPR, may have a subtler effect on peptide selection, but this remains to be addressed. Additional topics, such as recognition of peptides of length greater than 15 amino acids and C-terminal extensions of the peptide outside the MHC-I groove should be examined in future studies.

Based on our results, we propose a model for how a longer loop in the TAPBPR sequence may contribute in shaping the displayed peptide repertoire on MHC-I molecules. The G24-R36 loop serves as a trap, allowing specific peptides from the cellular pool to associate with MHC-I/TAPBPR complex with a binding affinity in the low micromolar range (relative to high μM, as observed for loop deletion mutants) **(Fig. 6A)**. By lowering the affinity requirements for binding across the peptide pool, TAPBPR can function as an auxiliary loader for peptides of low and moderate affinity, under conditions of high peptide concentration **(Fig. 6B)**. Local peptide concentrations will be highest in the vicinity of the TAP transporter, which acts as a hub around which PLC components associate. Tapasin lacks a long loop, but is tethered to the TAP transporter using a conserved Lys/Asp transmembrane salt-bridge (36) and therefore can still load cognate peptide ligands efficiently due to the increased local concentration **(Fig. 6C)**. In contrast, TAPBPR has lost the transmembrane Lysine and is not tethered to TAP, but instead utilizes a long loop acting as a kinetic trap to load peptides on empty or suboptimally loaded MHC-I molecules that have escaped the PLC. **(Fig. 6C)**. Irreversible peptide expulsion from the pMHC-I by a chaperone “scoop” is counterproductive for editing function, where the chaperone must transiently sense whether a pMHC-I complex is stable without release of bound peptide. By comparison, a peptide trap is best suited to editing function, whereby the loop acts as a lid over the MHC-I groove to slow the kinetics of peptide dissociation. The longer loop of TAPBPR is matched to its editing function in the Golgi where peptide concentrations will be low and only pMHC-I complexes with suboptimal peptides are to be disassembled, while tapasin has a short loop to favor competition between peptide substrates where they are in abundance surrounding the PLC **(Fig. 6C)**. In summary, our work offers a paradigm of how two closely related molecular chaperones have developed subtle variations of both static and dynamic conformational elements around a common structural theme, to achieve unique functions that are highly adapted to their specific cellular compartments.

**Figure 6.**
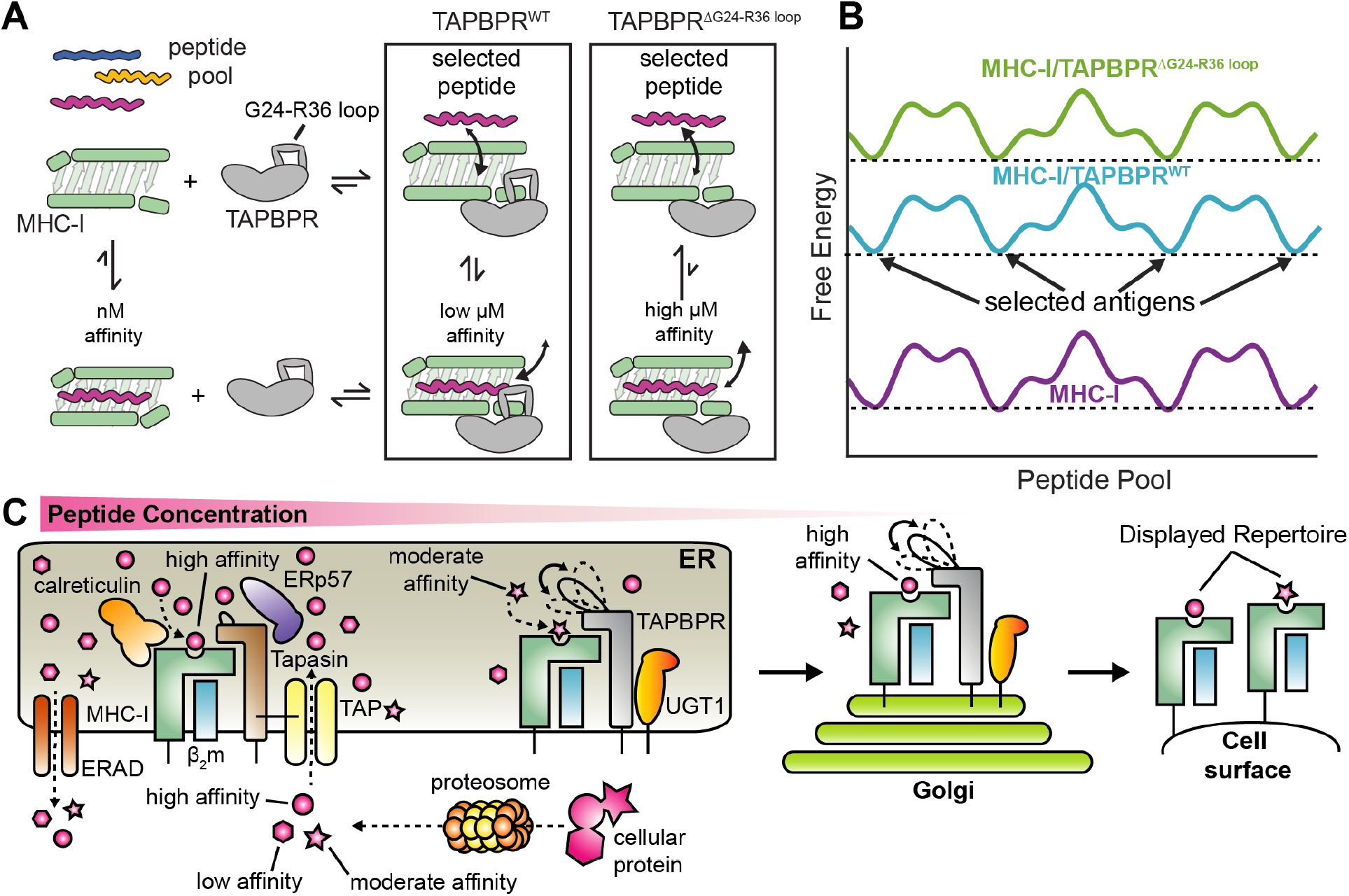
Proposed model of auxiliary peptide editing functions of TAPBPR. **(A)** Schematic of how the TAPBPR G24-R36 loop influences the peptide exchange cycle. The G24-R36 loop serves as a kinetic trap, promoting peptide binding on the MHC-I/TAPBPR complex with low μM affinity. When TAPBPR is mutated to have a shorter loop, the affinity of peptides is reduced to high μM range. **(B)** Conceptual example of the free energy landscape across the cellular peptide pool. The TAPBPR G24-R36 loop shapes the selected peptide repertoire by stabilizing peptide binding across the cellular pool, functioning as an auxiliary loading chaperone for peptides of low and moderate affinity. **(C)** In the ER, peptide selection is governed both by the effective concentration of peptides and the absence (in tapasin) or the presence of the kinetic trap (in TAPBPR). Tapasin is tethered to the TAP transporter and because it is restricted to an environment of high peptide concentration and has a shorter loop, it primarily loads high affinity peptides. In contrast, TAPBPR is not tethered to TAP and because it functions in environments of lower peptide concentration, it employs a trap to load both high and moderate affinity peptides and/or to minimize dissociation of peptides during transient TAPBPR/pMHC-I interactions. This process results in a diverse peptide repertoire being displayed by MHC-I molecules on the cell surface.

## Supporting information

Supplemental Methods, Figure S1 to S10, SI References

## Materials and Methods

Specific details about protein constructs, cell culture, flow cytometry, functional assays, immunoblots, yeast-display, deep-sequencing, recombinant protein expression and purification, ITC, NMR, FP experiments, Rosetta modeling and MD simulations, are outlined in detail in the SI Appendix.

## Reagent and Data Availability

Plasmids are deposited with Addgene. All Illumina sequencing data is deposited with GEO under series accession numbers GSE147137, GSE126206 and GSE118568. NMR assignments have been deposited into the Biological Magnetic Resonance Data Bank (http://www.bmrb.wisc.edu) under accession numbers 28107 and 28108.

## Author contributions

A.C.M., C.A.D., E.P., and N.G.S. designed research. A.C.M., C.A.D., N.A., and E.P. performed research. G.I.M., S.A.O., D.M. and N.A. contributed new reagents/analytic tools. A.C.M., C.A.D., E.P., and N.G.S. wrote the manuscript.

## Acknowledgements

We are grateful to Kannan Natarajan and David Margulies (NIH) for critical comments on our manuscript, and for providing the insect cell lines and DNA constructs for TAPBPR^WT^ and TAPBPR^ΔG24-R36^ protein expression. We thank Arne Schön (Johns Hopkins) for helpful comments on ITC experiments. E.P. was supported through NIAID (5R01AI129719). N.G.S. was supported through NIAID (5R01AI143997) NIGMS (5R35GM125034) and High End Instrumentation (HIE) Grant S10OD018455, which funded the 800 MHz NMR spectrometer at UCSC. Flow cytometry and Illumina sequencing were supported by the UIUC Roy J. Carver Biotechnology Center.

## References

1. J. S. Blum, P. A. Wearsch, P. Cresswell, Pathways of antigen processing. Annu. Rev. Immunol. 31, 443–473 (2013).

2. J. Rossjohn, et al., T cell antigen receptor recognition of antigen-presenting molecules. Annu. Rev. Immunol. 33, 169–200 (2015).

3. A. van Hateren, A. Bailey, T. Elliott, Recent advances in Major Histocompatibility Complex (MHC) class I antigen presentation: Plastic MHC molecules and TAPBPR-mediated quality control. F1000Research 6, 158 (2017).

4. C. Hermann, J. Trowsdale, L. H. Boyle, TAPBPR: a new player in the MHC class I presentation pathway. Tissue Antigens 85, 155–166 (2015).

5. A. C. McShan, et al., Molecular determinants of chaperone interactions on MHC-I for folding and antigen repertoire selection. Proc. Natl. Acad. Sci. U. S. A. 116, 25602–25613 (2019).

6. C. Hermann, et al., TAPBPR alters MHC class I peptide presentation by functioning as a peptide exchange catalyst. eLife 4 (2015).

7. G. Fleischmann, et al., Mechanistic Basis for Epitope Proofreading in the Peptide-Loading Complex. J. Immunol. Baltim. Md 1950 195, 4503–4513 (2015).

8. G. I. Morozov, et al., Interaction of TAPBPR, a tapasin homolog, with MHC-I molecules promotes peptide editing. Proc. Natl. Acad. Sci. U. S. A. 113, E1006–1015 (2016).

9. A. Neerincx, L. H. Boyle, Properties of the tapasin homologue TAPBPR. Curr. Opin. Immunol. 46, 97–102 (2017).

10. Y. Shionoya, et al., Loss of tapasin in human lung and colon cancer cells and escape from tumor-associated antigen-specific CTL recognition. Oncoimmunology 6, e1274476 (2017).

11. Q.-R. Chen, Y. Hu, C. Yan, K. Buetow, D. Meerzaman, Systematic Genetic Analysis Identifies Cis-eQTL Target Genes Associated with Glioblastoma Patient Survival. PLOS ONE 9, e105393 (2014).

12. B. Park, et al., Human cytomegalovirus inhibits tapasin-dependent peptide loading and optimization of the MHC class I peptide cargo for immune evasion. Immunity 20, 71–85 (2004).

13. V. Montserrat, B. Galocha, M. Marcilla, M. Vázquez, J. A. López de Castro, HLA-B*2704, an allotype associated with ankylosing spondylitis, is critically dependent on transporter associated with antigen processing and relatively independent of tapasin and immunoproteasome for maturation, surface expression, and T cell recognition: relationship to B*2705 and B*2706. J. Immunol. Baltim. Md 1950 177, 7015–7023 (2006).

14. J. H. Lee, et al., Further Examination of the Candidate Genes in Chromosome 12p13 Locus for Late-Onset Alzheimer Disease. Neurogenetics 9, 127–138 (2008).

15. C. Thuring, et al., HLA class I is most tightly linked to levels of tapasin compared with other antigen-processing proteins in glioblastoma. Br. J. Cancer 113, 1640 (2015).

16. T. Cabrera, M. A. López-Nevot, J. J. Gaforio, F. Ruiz-Cabello, F. Garrido, Analysis of HLA expression in human tumor tissues. Cancer Immunol. Immunother. CII 52, 1–9 (2003).

17. C. M. Cabrera, M.-A. López-Nevot, P. Jiménez, F. Garrido, Involvement of the chaperone tapasin in HLA-B44 allelic losses in colorectal tumors. Int. J. Cancer 113, 611–618 (2005).

18. J. Jiang, et al., Crystal structure of a TAPBPR–MHC I complex reveals the mechanism of peptide editing in antigen presentation. Science 358, 1064–1068 (2017).

19. C. Thomas, R. Tampé, Structure of the TAPBPR–MHC I complex defines the mechanism of peptide loading and editing. Science 358, 1060–1064 (2017).

20. A. Blees, et al., Structure of the human MHC-I peptide-loading complex. Nature 551, 525 (2017).

21. I. Hafstrand, et al., Successive crystal structure snapshots suggest the basis for MHC class I peptide loading and editing by tapasin. Proc. Natl. Acad. Sci. U. S. A. 116, 5055–5060 (2019).

22. L. Sagert, F. Hennig, C. Thomas, R. Tampé, A loop structure allows TAPBPR to exert its dual function as MHC I chaperone and peptide editor. eLife 9, e55326 (2020).

23. F. T. Ilca, et al., TAPBPR mediates peptide dissociation from MHC class I using a leucine lever. eLife 7 (2019).

24. A. Leaver-Fay, et al., ROSETTA3: an object-oriented software suite for the simulation and design of macromolecules. Methods Enzymol. 487, 545–574 (2011).

25. A. C. McShan, et al., Molecular determinants of chaperone interactions on MHC-I for folding and antigen repertoire selection. Proc. Natl. Acad. Sci. U. S. A. 116, 25602–25613 (2019).

26. E. Procko, et al., Computational design of a protein-based enzyme inhibitor. J. Mol. Biol. 425, 3563–3575 (2013).

27. J. Karanicolas, et al., A De Novo Protein Binding Pair By Computational Design and Directed Evolution. Mol. Cell 42, 250–260 (2011).

28. A. C. McShan, et al., Peptide exchange on MHC-I by TAPBPR is driven by a negative allostery release cycle. Nat. Chem. Biol. 14, 811–820 (2018).

29. A. R. Khan, B. M. Baker, P. Ghosh, W. E. Biddison, D. C. Wiley, The structure and stability of an HLA-A*0201/octameric tax peptide complex with an empty conserved peptide-N-terminal binding site. J. Immunol. Baltim. Md 1950 164, 6398–6405 (2000).

30. C. Hassan, et al., Naturally processed non-canonical HLA-A*02:01 presented peptides. J. Biol. Chem. 290, 2593–2603 (2015).

31. F. E. Tynan, et al., High resolution structures of highly bulged viral epitopes bound to major histocompatibility complex class I. Implications for T-cell receptor engagement and T-cell immunodominance. J. Biol. Chem. 280, 23900–23909 (2005).

32. T. Trolle, et al., The length distribution of class I restricted T cell epitopes is determined by both peptide supply and MHC allele specific binding preference. J. Immunol. Baltim. Md 1950 196, 1480–1487 (2016).

33. C. Thomas, R. Tampé, Structure of the TAPBPR-MHC I complex defines the mechanism of peptide loading and editing. Science, eaao6001 (2017).

34. J. Jiang, et al., Crystal structure of a TAPBPR-MHC-I complex reveals the mechanism of peptide editing in antigen presentation. Science, eaao5154 (2017).

35. F. T. Ilca, L. Z. Drexhage, G. Brewin, S. Peacock, L. H. Boyle, Distinct Polymorphisms in HLA Class I Molecules Govern Their Susceptibility to Peptide Editing by TAPBPR. Cell Rep. 29, 1621–1632.e3 (2019).

36. A. Blees, et al., Assembly of the MHC I peptide-loading complex determined by a conserved ionic lock-switch. Sci. Rep. 5, 17341 (2015).

