## Supplemental Methods, Figure S1 to S10, SI References for "TAPBPR Promotes Antigen Loading on MHC-I Molecules Using a Peptide Trap"

**This PDF file includes:**

Supplemental Methods

Figure S1 to S10

SI References

### Supplemental Methods

#### Human Cell Expression Constructs

For expression in human cells, cDNAs were cloned into the NheI-XhoI sites of pCEP4 (Invitrogen) with a consensus Kozak sequence. A synthetic codon-optimized sequence of human tapasin (isoform 1 a.a. 21-448; GenBank Acc. No. NP_003181) was synthesized (IDT) with a canonical signal peptide and N-terminal FLAG epitope tag. The cloning of TAPBPR is previously described(1). TAPBPR-CT was constructed by PCR-based fragment assembly: the final sequence comprises a canonical signal peptide, a FLAG tag, TAPBPR residues 22-405 and tapasin residues 406-448. Both targeted mutations and generation of the TAPBPR-CT libraries using oligos with degenerate NNK codons were by overlap extension PCR(2).

#### Yeast Expression Constructs

For yeast surface display, the extracellular domain of TAPBPR (a.a. 22-405) was cloned into the NheI-XhoI sites of pETCON(3), fusing Aga2p and a c-myc tag to the TAPBPR N- and C-termini for surface display and detection, respectively. To generate the SSM library on the 24-35 loop, the TAPBPR insert was amplified using oligos with degenerate NNK codons by overlap extension PCR. The pooled PCR product was mixed with linearized pETCON (cut NdeI/XhoI) and electroporated in to yeast, thereby using natural homologous recombination pathways to fuse the TAPBPR insert in to the pETCON backbone.

#### Human Cell Culture

Expi293F cells were cultured in suspension at 37 °C, 8% $CO_2$, 125 rpm using Expi293 Expression Media (ThermoFisher). To generate the tapasin knockout line, cells were co-transfected with plasmids(4) encoding human codon-optimized Streptococcus pyogenes Cas9 and two guide RNAs targeting the tapasin gene immediately following the signal peptide (target sequences 5'-GACCCGCGGTGATCGAGTGTTGG-3' and 5'-AACCAACACTCGATCACCGCGGG-3'). After a week, cells with reduced surface MHC-I following staining with anti-HLA-A2-PE (1/200 dilution, clone BB7.2, Biolegend) were enriched by sorting on a BD FACSAria II. Genomic DNA was purified (DNeasy Blood and Tissue Kit, Qiagen), and the targeted region of the tapasin gene was PCR amplified and sequenced on an Illumina MiSeq. Based on analysis with CRISPR-GA(5), >99 % of tapasin gene copies in the polyclonal knockout line had indel mutations (GEO sample ID GSM3593598), whereas copies of the tapasin gene were mostly wild type in the parental line (GEO sample ID GSM3593597). To test targeted mutants of TAPBPR or tapasin, Expi293F cells (wild type or tapasin-KO) were transfected using ExpiFectamine (Life Technologies) with 500 ng plasmid DNA per ml of cells at a density of $2 \times 10^6$ / ml. Cells were analyzed 24–26 h post-transfection. For sorting libraries, tapasin-KO Expi293F cells ($2 \times 10^6$ / ml) were transfected with 1 ng SSM library plasmid diluted with 1500 ng carrier plasmid (pCEP4-ΔCMV(6)) using ExpiFectamine, and the medium was replaced after 2 hours. Cells were sorted 24-25 h post-transfection.

#### Flow Cytometry of Expi293F Cells

To assess surface HLA-A2 expression, cells were stained on ice for 30 minutes with anti-HLA-A2 PE (clone BB7.2, BioLegend) diluted 1:200 in PBS supplemented with 0.2% bovine serum albumin (PBS-BSA) and then washed twice with PBS-BSA. Fluorescence was measured using a BD LSRII and results were analyzed using FCS Express 6.

**Deep Mutational Scan of TAPBPR-CT for Chaperone Activity in Human Cells**
Tapasin-KO Expi293F cells expressing the TAPBPR-CT library were stained with 1:200 anti-HLA-A2 PE in PBS-BSA for 30 minutes at 4 °C and washed twice with PBS-BSA. After gating for the main population by foward-side scatter and excluding dead cells based on DAPI uptake, the top 0.5% of cells for PE fluorescence were collected using a BD FACSAria II. RNA was extracted from sorted cells (GeneJet RNA purification kit, ThermoFisher), first strand cDNA was synthesized using Accuscript (Agilent) primed with the EBV reverse sequencing primer, which anneals to the 3' UTR of pCEP4-encoded transcripts. DNA fragments covering mutated regions were PCR amplified in two rounds to append sequences complementary to Illumina sequencing primers and add experiment-specific barcodes and Illumina adaptamers. Products were sequenced on a NovaSeq 6000 and analyzed with Enrich(7). Scripts for analysis are included with the GEO submission.

**Immunoblots**
Cells were pelleted, lysed in reducing SDS load dye and sonicated. Proteins were separated by electrophoresis on 10% SDS polyacrylamide gels and transferred to PVDF. Membranes were blocked with 3% BSA in Tris-buffered saline supplemented with 0.1% tween 20 (TBS-T) for 30 mins, followed by staining with 1:2000 anti-FLAG (clone M2)-AP (Sigma-Aldrich) for 30 mins, washed 5 times with TBS-T, and developed using 1-Step NBT-BCIP (ThermoFisher).

**Yeast Display**
Saccharomyces cerevisiae strain EBY100 were grown in YPAD medium. To test targeted mutants, lithium acetate/polyethylene glycol 3350-treated competent yeast were heat shocked to promote plasmid uptake. For generating the TAPBPR 24-35 loop library, yeast were electroporated. Transformed yeast were selected in SDCAA medium (2% w/v glucose, 0.67% w/v yeast nitrogen base, 0.5% w/v casamino acids, 0.1 M sodium phosphate pH 6.6) at 30 °C for 1-2 days, before induction in SGCAA (in which glucose is replaced with galactose) at OD(600 nm) = 0.5 for 2 days at 24 °C. Induced EBY100 were washed with PBS-BSA and incubated with APC-conjugated MHC-I tetramers (concentrations are indicated in figure legends) and FITC-conjugated chicken anti-c-myc (1/100 dilution, Immunology Consultants Laboratory) for 40 minutes at 24°C, 215 rpm. Cells were washed twice with cold PBS-BSA before being resuspended for analysis on a BD Accuri C6 or sorted on a BD FACSAria II.

**Deep Mutational Scan of TAPBPR 24-35 Loop using Yeast Display**
Generation of the TAPBPR library and yeast preparation are described above. Naive and sorted yeast cultures were lysed with 125 U/ml Zymolase (37˚C, 5 hr) and plasmid DNA was purified using a Zymoprep kit (Zymo Research). The mutated region of TAPBPR was PCR amplified in two stages. A first round of PCR used primers that added sequences complementary for Illumina sequencing primers. A second round of PCR added end sequences for annealing to the Illumina flow cell and included 6 bp barcodes for unique sample identification. Amplicons were sequenced on a Illumina HiSeq 4000 and data were analyzed with Enrich(7). Scripts for analysis are included in the GEO submission.

**Protein Expression and Purification.** DNA plasmid constructs encoding the luminal domain of human HLA-A*02:01 (heavy chain) and h$\beta_2$m (light chain) were generously provided by the NIH tetramer facility and transformed into BL21(DE3) *Escherichia coli* (New England Biolabs). DNA

Plasmid construct encoding the luminal domain of mouse H2-D$^d$ molecule was generously provided by Kannan Natarajan, NIH. MHC-I heavy and light chain molecules were individually expressed in Luria-Broth, extracted from inclusion bodies, and refolded *in vitro* together with peptide at 4°C as previously described(8). Peptides used in this study were prepared by chemical synthesis (Biopeptik Inc, Malvern, USA or GenScript, Piscataway, USA). Peptides sequences include: P18-I10 (RGPGRAFVTI), TAX8 (LFGYPVYV), TAX9 (LLFGYPVYV), TAX10 (LLFGGYPVYV), TAX11 (LLFGGGYPVYV), TAX12 (LLFGGGGYPVYV) and KLL15 (KLLEIPDPDKNWATL). The UV-labile conditional ligands photoP18-I10 (RGPGRAF*J*TI) and photoFluM1 (KILGFVF*J*V), where *J* = 3-amino-3-(2-nitrophenyl)-propionic acid were refolded with H2-D$^d$/h$\beta_2$m and HLA-A*02:01/h$\beta_2$m, respectively, under dark conditions (9). Purification of pMHC-I complexes was performed by size-exclusion chromatography (SEC) with a HiLoad 16/600 Superdex 75 pg column at 1 mL/min with running buffer (150 mM NaCl, 25 mM Tris, pH 8). The luminal domains of various TAPBPR forms used in this study were expressed using a Drosophila S2 cell expression system and purified as previously described (10). The TAPBPR$^{\Delta ALAS}$ construct was prepared by PCR using through deletion of A29-S32 using forward primer: 5' – AAG GAC GGT GCG CAC CGT GGA AGT GAG GAC AGG GCA AGG GCC – 3' and reverse primer: 5' – GGC CCT TGC CCT GTC CTC ACT TCC ACG GTG CGC ACC GTC CTT – 3', and confirmed by DNA sequencing. The TAPBPR$^{WT}$ and TAPBPR$^{\Delta G24\text{-}R36}$ constructs were generously provided by Kannan Natarajan, NIH. DNA transfection of the TAPBPR constructs into Drosophila S2 cells was performed using standard protocols with X-tremeGENE 9 (Sigma-Aldrich). Following expression and purification all proteins were exhaustively buffer exchanged into 50 mM NaCl, 20 mM sodium phosphate pH 7.2.

**Sequence alignment.** Sequence alignment was performed between *Homo sapiens* (Hs, human) TAPBPR (UniProtID: Q9BX59) and tapasin (UniProtID: O15533), *Mus musculus* (Mm, mouse) TAPBPR (UniProtID: Q8VD31) and tapasin (UniProtID: Q9R233), and *Rattus norvegicus* (Rn, rat) TAPBPR (UniProtID: D4A6L1) and tapasin (UniProtID: Q99JC6) using ClustalOmega.

***Rosetta* Modeling.** *Rosetta* modeling of MHC-I/TAPBPR complexes and the TAPBPR G24-R36 loop was performed using either *RosettaCCD* (11), *RosettaKIC* (12), or *RosettaCM* (13) using template X-ray structure PDB ID 5WER or PDB ID 5OPI. In each case the sequence for H2-D$^d$ or H2-D$^b$ was replaced with HLA-A*02:01 using *Rosetta*'s partial_thread application. Fragment files for loop modeling were generated using the Robetta server (http://robetta.bakerlab.org/). The lowest energy structures were selected from a total of 1,000 models calculated for each protocol. *Rosetta* modeling of TAX10, TAX11 or TAX12 peptides in complex with HLA-A*02:01 was performed using *RosettaCM* against PDB ID 1HHH, 5D9S, and 4JQX, respectively.

**Molecular dynamics simulations**. All-atom molecular dynamics (MD) simulations in explicit solvent were carried out as previously described in GROMACS version 2019.2 using an AMBER99SB-ILDN protein force field and TPI3P water model (14). The input structure was the lowest energy *RosettaCM* model of peptide-deficient HLA-A*02:01/h$\beta_2$m/TAPBPR built from template PDB ID 5OPI. LINCS and SETTLE constraint algorithms were used to constrain protein and water molecules, respectively. An integration time step of 4 fsec was used with coordinates output every 10 psec. Short range interactions were treated with a Verlet cut-off scheme with 10 Å electrostatic and van der Walls cutoffs and long-range electrostatics were treated with the PME method with a grid spacing of 1.2 Å and cubic interpolation. Periodic dodecahedron boundaries were used. The thermodynamic ensemble was nPT where temperature was kept constant at 300K by a V-rescale modified thermostat with 0.1 psec time constant and pressure was kept constant at 1 bar pressure using an isotropic Berendsen barostat. The system was solvated to overall neutral charge and contained $Na^+$ and $Cl^-$ ions to yield physiological concentration of 0.15 M. Following 500 steps of steepest-descent energy minimization, initial velocities were generated at 65 K with linear heating up to 300 K over 2 nsec. Trajectories were acquired for 200 nsec. Structures were extracted after the final 200 nsec simulations using GROMACS.

**Differential Scanning Fluorimetry.** DSF experiments were performed on an Applied Biosystems ViiA 7 qPCR machine with excitation and emission wavelengths set to 470 nm and 569 nm with proteins in buffer of 50 mM NaCl, 20 mM sodium phosphate pH 7.2. Experiments were conducted in triplicate in MicroAmp Fast 96-well plates with 50 μL total volume containing final concentrations of 7 μM protein and 10× SYPRO orange dye (ThermoFisher). Temperature was incrementally increased at a scan rate of 1°C/min between 25°C and 95°C. Data analysis and fitting were performed in GraphPad Prism v7.

**Circular Dichroism.** Far-UV CD spectra were acquired using a JASCO J-815 Spectropolarimeter. CD spectra of wild-type TAPBPR, $TAPBPR^{\Delta ALAS}$ and $TAPBPR^{\Delta G24\text{-}R36}$ were acquired using 0.05 mg/mL protein in 2 mL of 50 mM NaCl, 20 mM sodium phosphate pH 7.2 in a quartz cuvette. A buffer blank was acquired and subtracted from each CD spectra. CD spectra were acquired from 190 to 260 nm in triplicate at 25°C with a scan rate of 50 nm/minute. The experimental CD values of ellipticity (mdeg) were converted to molar ellipticity ($\theta$ = deg $cm^2$/dmol).

**NMR Spectroscopy.** Samples for NMR were prepared using either $ILV^{proS}$ (Ile $^{13}C\delta 1$, Leu $^{113}C\delta 2$, Val $^{13}C\gamma 2$) or AILV methyl (Ala $^{13}C\beta$, Ile $^{13}C\delta 1$, Leu $^{13}C\delta 1/^{13}C\delta 2$, Val $^{13}C\gamma 1/^{13}C\gamma 2$) isotopic labeling at the HLA-A*02:01 heavy chain against a $^{12}C/^{2}H/^{15}N$ background. Bound peptide, $h\beta_2m$ or TAPBPR were fully protonated. NMR methyl resonance assignments of free and TAPBPR bound states of HLA-A*02:01 were reported previously by our group (1). Peptide-deficient HLA-A*02:01/$h\beta_2m$/TAPBPR complexes were prepared as described above in the section "Preparation of empty MHC-I/TAPBPR complexes" where HLA-A*02:01 was $ILV^{proS}$ or AILV methyl labeled. The resulting purified HLA-A*02:01/$h\beta_2m$/$TAPBPR^{WT}$, HLA-A*02:01/$h\beta_2m$/$TAPBPR^{\Delta ALAS}$, or HLA-A*02:01/$h\beta_2m$/$TAPBPR^{\Delta G24\text{-}R36}$ complexes were exhaustively dialyzed into NMR buffer (50 mM NaCl, 20 mM sodium phosphate pH 7.2, 5% $D_2O$) and concentrated to ~ 80 μM. Two-dimensional $^{1}H$-$^{13}C$ methyl SOFAST HMQC experiments (15) were recorded at 25ºC at a $^{1}H$ field strength of 800 MHz. A total number of 320 scans were used with a 0.2 sec recycle delay (d1) and acquisition times of 30 msec in the $^{13}C$ dimension. Data were processed with 4 Hz and 10 Hz Lorentzian line broadening in the direct and indirect dimensions. Chemical shift deviations (CSD, p.p.m.) were determined between TAX9/HLA-A*02:01/$h\beta_2m$ and TAPBPR bound HLA-A*02:01/$h\beta_2m$ using the equation $\Delta\delta^{CH3}=[1/2(\Delta\delta_H^{2}+\Delta\delta_C^{2}/4)]^{1/2}$ for each methyl resonance. All NMR data were processed with NMRPipe (16) and analyzed using NMRFAM-SPARKY (17).

**Isothermal Titration Calorimetry.** ITC was performed using a MicroCal VP-ITC system (Malvern Panalytical). All proteins were exhaustively dialyzed into the buffer (50 mM NaCl, 20 mM sodium phosphate pH 7.2) filtered through a 0.22 μm PES membrane. ITC experiments to probe the $K_{D2}$ step of the peptide exchange cycle were performed under substoichiometric conditions in the absence of excess TAPBPR to allow for dissociation of HLA-A*02:01 from TAPBPR in the presence of incoming peptide. Syringe containing ~50 to 100 μM peptide was titrated into a calorimetry cell containing ~15 μM purified peptide-deficient HLA-A*02:01/h$\beta_2$m/TAPBPR complex. ITC experiments to probe the $K_{D3}$ step of the peptide exchange cycle were performed by titration of ~ 150 to 200 μM purified pMHC-I into a calorimetry cell containing ~15 μM TAPBPR and 1 mM excess peptide. ITC experiments to probe the $K_{D4}$ step of the peptide exchange cycle were performed under stoichiometric conditions in the presence of excess TAPBPR to minimize dissociation of TAPBPR from the pMHC-I/TAPBPR complex. Syringe containing ~ 150 to 200 μM peptide was titrated into a calorimetry cell containing ~15 μM peptide-deficient HLA-A*02:01/h$\beta_2$m/TAPBPR complex and 50 μM excess TAPBPR. In all ITC experiments injection volumes were 10 μL performed for a duration of 10 sec and spaced 220 sec apart to allow for a complete return to baseline. Data was subtracted from a control experiments. Data were processed and analyzed with Origin software. Isotherms were fit using a one-site ITC binding model. The first data point was excluded from analysis.

**Fluorescence Polarization.** FP was performed using a modified TAX9 peptide labeled with fluorescent TAMRA dye ($K^{TAMRA}$LFGYPVYV, herein called TAMRA-TAX9) (Biopeptik Inc, Malvern, USA). Peptide-deficient HLA-A*02:01/h$\beta_2$m/TAPBPR complexes for FP were prepared by UV-irradiation at 365 nm for 1 hour of photoFluM1/HLA-A*02:01/h$\beta_2$m/TAPBPR complexes followed by purification, as described in the section "Preparation of empty MHC-I/TAPBPR complexes". FP experiments to probe the $K_{D2}$ step of the peptide exchange cycle (defined as $IC_{50\ 2}$ in the FP competition experiments) were performed under substoichiometric conditions in the absence of excess TAPBPR to allow for dissociation of HLA-A*02:01/h$\beta_2$m from TAPBPR in the presence of incoming peptide. Graded concentrations (0 nM, 2.5 nM, 5 nM, 10 nM, 25 nM, 50 nM, 100 nM, 500 nM, 1000 nM, 2000 nM, 3000 nM, 4000 nM and 5000 nM) of TAX8, TAX9, TAX10, TAX11, TAX12, or KLL15 were added to a mixture of 1 nM TAMRA-TAX9 and either 50 nM of peptide-deficient HLA-A*02:01/h$\beta_2$m/$TAPBPR^{WT}$, peptide-deficient HLA-A*02:01/h$\beta_2$m/$TAPBPR^{\Delta ALAS}$, or peptide-deficient HLA-A*02:01/h$\beta_2$m/$TAPBPR^{\Delta G24\text{-}R36}$. FP experiments to probe the $K_{D3}$ step of the peptide exchange cycle were performed by titration of graded concentrations (0 μM, 0.1 μM, 2 μM, 4 μM, 10 μM, 30 μM, 50 μM and 80 μM) of $TAPBPR^{WT}$, $TAPBPR^{\Delta ALAS}$ or $TAPBPR^{\Delta G24\text{-}R36}$ into 1 nM TAMRA-TAX9 and 50 nM TAX8/HLA-A*02:01/h$\beta_2$m. FP experiments to probe the $K_{D4}$ step of the peptide exchange cycle (defined as $IC_{50\ 4}$ in the FP competition experiments) were performed under stoichiometric conditions in the presence of excess $TAPBPR^{WT}$, $TAPBPR^{\Delta ALAS}$ or $TAPBPR^{\Delta G24\text{-}R36}$ to minimize dissociation of TAPBPR from the pMHC-I/TAPBPR complex. Graded concentrations (0 nM, 10 nM, 50 nM, 250 nM, 500 nM, 1000 nM, 2000 nM, 3000 nM, 4000 nM, 6000 nM, 8000 nM, 10,000 nM, 50,000 nM, 100,000 nM) of TAX8, TAX9, TAX10, TAX11, TAX12, or KLL15 were added to a mixture of 1 nM TAMRA-TAX9 and either 50 nM of peptide-deficient HLA-A*02:01/h$\beta_2$m/$TAPBPR^{WT}$, peptide-deficient HLA-A*02:01/h$\beta_2$m/$TAPBPR^{\Delta ALAS}$, or peptide-deficient HLA-A*02:01/h$\beta_2$m/$TAPBPR^{\Delta G24\text{-}R36}$ together with 1 μM of their respective free $TAPBPR^{WT}$, $TAPBPR^{\Delta ALAS}$ or $TAPBPR^{\Delta G24\text{-}R36}$. Each experiment was performed in a volume of 140 μL and loaded onto a black 96-well polystyrene assay plate (Costar 3915). FA data was recorded via a Perkin-Elmer Envision 2103 plate reader with excitation filter $\lambda_{ex}$ = 531 nm and emission filter $\lambda_{em}$ = 595 nm with measurement height 4.3, excitation light 100, G-factor 1.36 and a total of 100 flashes. In each of the above experiments, the average of FP of after incubation for 95-105 minutes 25°C was plotted as a function of the $\log_{10}$ of excess peptide. Each experiment was performed in triplicate and is representative of at least two independent experiments. Experimental values were subtracted from background FA values obtained from incubation of TAMRA-TAX9 alone. All samples were prepared in matched buffer (50 mM NaCl, 20 mM sodium phosphate pH 7.2, 0.05% (v/v) tween-20). Data were fit using GraphPad Prism v7.

### Supplemental Figures

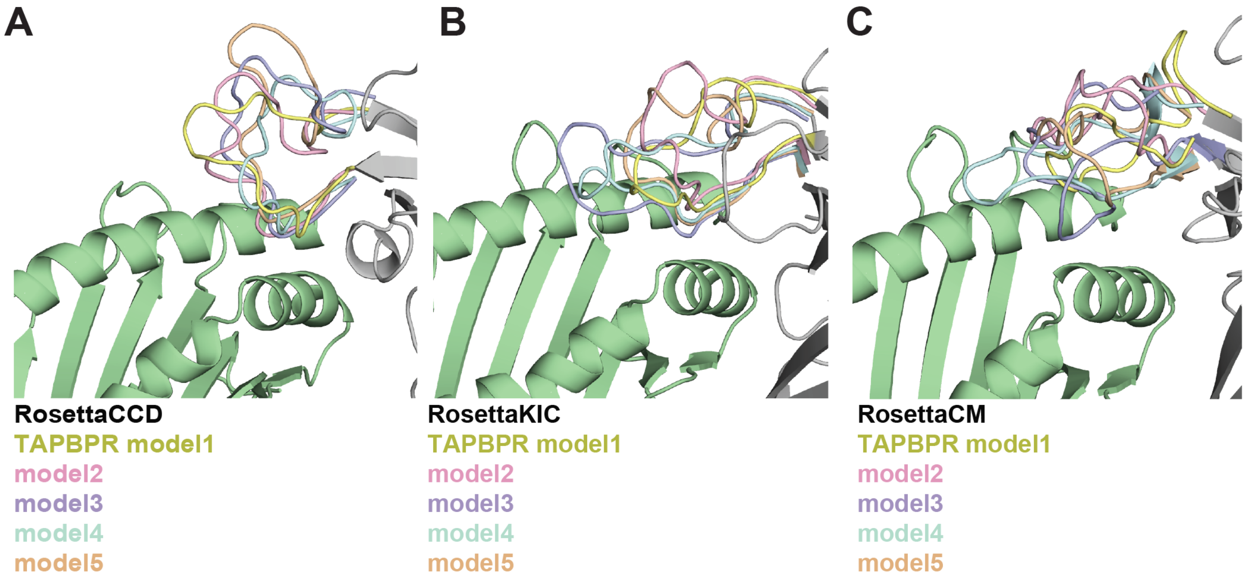


**Figure S1. *Rosetta* modeling of the TAPBPR G24-R36 loop.** Different *Rosetta* protocols are used to model the TAPBPR G24-R36 loop. **(A)** Cyclic coordinate descent (CCD), **(B)** kinematic closure (KIC), or **(C)** comparative modeling (CM). The template used was PDB ID 5WER where HLA-A*02:01 sequence was threaded onto H2-$D^d$ using the partial_thread application in *Rosetta*. HLA-A*02:01 is green; TAPBPR is gray. The five lowest energy models of the TAPBPR G24-R36 loop from a total of 1,000 calculated are shown in different colors.

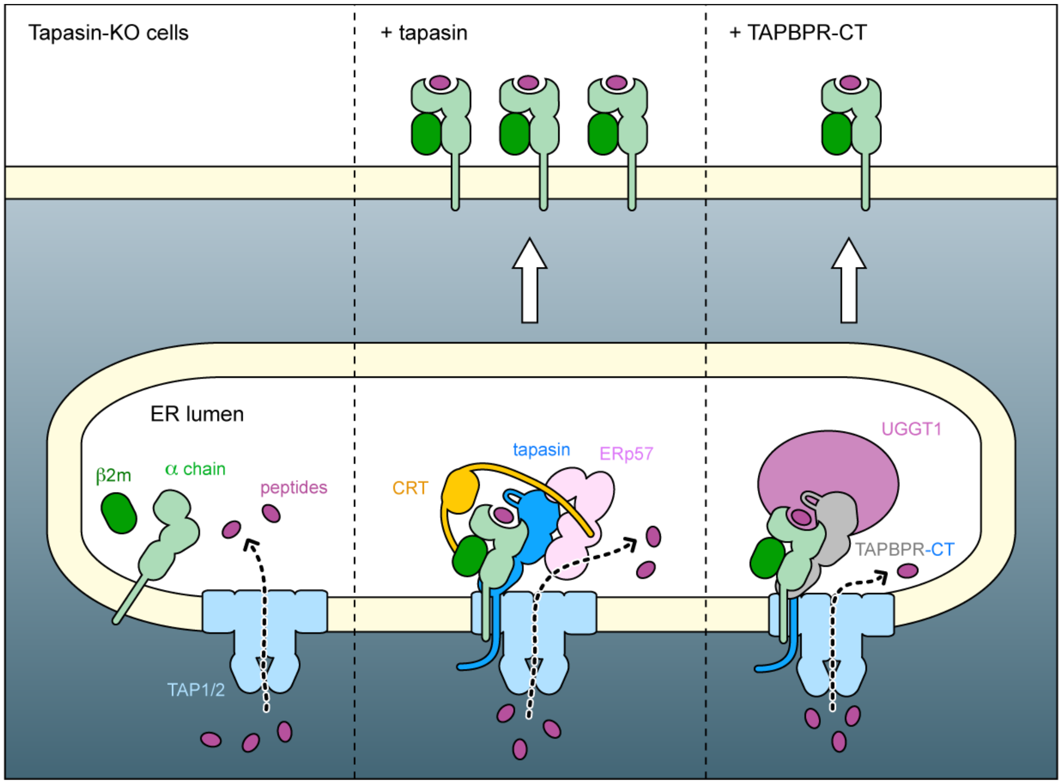


**Figure S2. Functional replacement of endogenous tapasin for HLA-A2 processing and surface trafficking.** (*Left*) In tapasin-KO cells, HLA-A2 is poorly processed and loaded, and is retained intracellularly. (*Center*) Expression of tapasin from a plasmid rescues formation of the peptide-loading complex (PLC). HLA-A2 is now efficiently loaded with peptides, folds, and traffics to the plasma membrane. (*Right*) Expression of a chimera between the TAPBPR extracellular domains and the tapasin transmembrane and cytosolic domains partially rescues HLA-A2 processing. We hypothesize that the tapasin C-terminal tail might help bring TAPBPR and its associated glucosyltransferase UGGT1 in to a PLC-like complex with the TAP1/2 transporter.

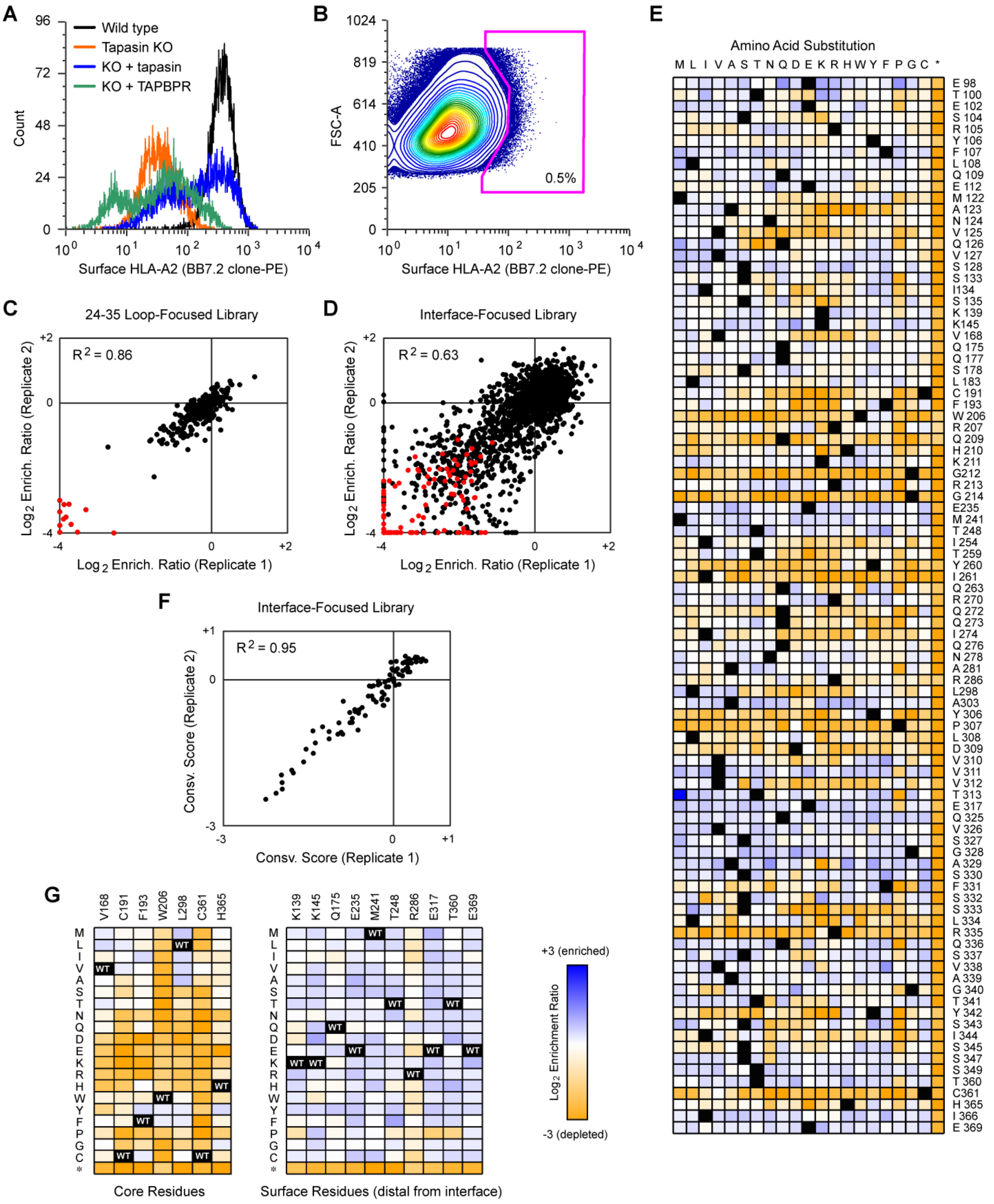


**Figure S3. Selection strategy and data quality for deep mutagenesis of TAPBPR. (A)** Tapasin was knocked out by targeting indel mutations to exon 2 in Expi293F cells using Cas9/CRISPR-based editing (confirmed by deep sequencing genomic DNA; data deposited in GEO). Surface HLA-A*02:01 expression was dramatically reduced in tapasin-KO cells (red) compared to wild type cells (black). Surface trafficking of HLA-A*02:01 was rescued by transfection with a plasmid encoding tapasin (blue), or partially rescued by over-expression of TAPBPR (green). When TAPBPR is over-expressed, a subset of cells also shows increased intracellular retention of HLA-A*02:01 (represented by the left-most peak in green). **(B)** Libraries of TAPBPR-CT variants were expressed in tapasin-KO Expi293F cells, and after first gating by scattering and viability, the 0.5 % of cells with the highest levels of endogenous HLA-A2 at the cell surface were collected (magenta gate). Under these transfection conditions, cells typically express no more than a single sequence variant, and most cells are negative. **(C)** Following selection of a TAPBPR-CT library for rescue of surface HLA-A2 expression in tapasin-KO cells, $\log_2$ enrichment ratios for nonsynonymous mutations (black) in the 24-35 loop cluster near the origin. Nonsense mutations (red) are depleted. Data from two independent experiments are plotted on the x- and y-axes. **(D)** Agreement of mutation enrichment ratios after two independent sorting experiments of a TAPBPR-CT library encoding 1,840 substitutions at 92 amino acid positions. Nonsynonymous mutations are black, nonsense mutations are red. **(E)** $\text{Log}_2$ enrichment ratios from D, are plotted as a heatmap, colored from orange (≤ -3, i.e. depleted/deleterious) to pale/white (0, i.e. neutral) to dark blue (≥ +3, i.e. enriched). Due to read length limits for Illumina sequencing, enrichment of the wild type sequence in the experiment is unknown and shown in black. Mutagenized TAPBPR positions are on the vertical axis, while amino acid substitutions are on the horizontal axis (*, stop codon). **(F)** From data in D, conservation scores are calculated from the mean of the $\log_2$ enrichment ratios for all amino acid substitutions at each residue position. Conservation scores are closely correlated between independent sorting experiments. TAPBPR residues with negative scores are tightly conserved for functional rescue of surface HLA-A*02:01. (G) Heatmaps colored as in E, showing that core TAPBPR residues are restricted to hyrophobics (*left*) while surface TAPBPR residues distal from the MHC-I interface are mutationally tolerant (*right*).

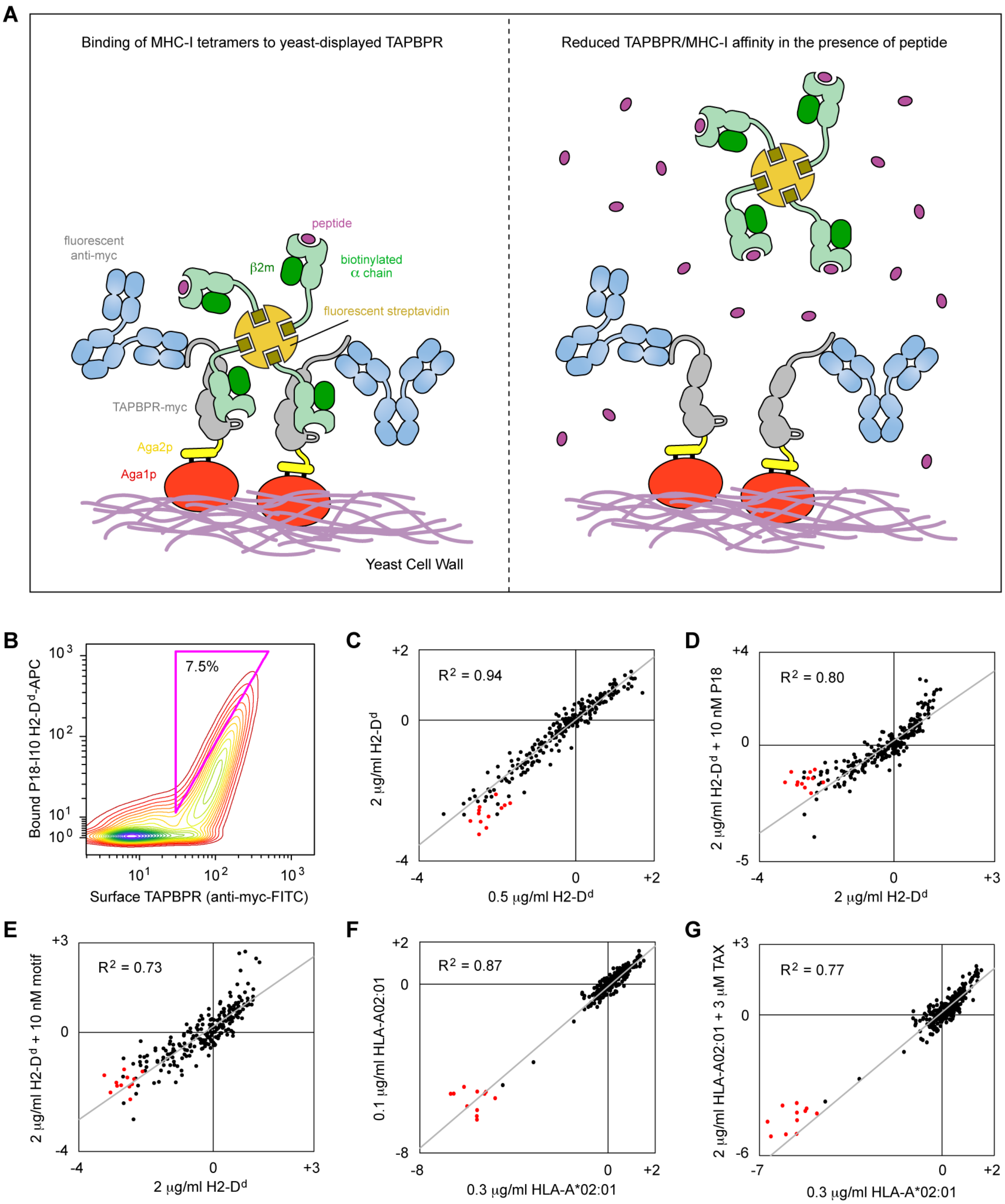


**Figure S4. Yeast display for assessing how TAPBPR sequence influences binding to folded MHC-I. (A)** The extracellular domains of TAPBPR are expressed on the surface of the yeast cell wall via an N-terminal fusion with Aga2p. A c-myc epitope tag at the C-terminus is used for fluorescence detection of expression levels with anti-myc antibodies. MHC-I binding is measured by flow cytometry after incubation of the yeast with fluorescent pMHC-I tetramers. TAPBPR binds MHC-I with highest affinity when it is peptide free, and the addition of peptide reduces MHC-I binding to TAPBPR-expressing yeast. **(B)** The 24-35 loop of TAPBPR was diversified by saturation mutagenesis. The yeast-displayed library was sorted under a variety of conditions for binding to tetramers of P18-I10-loaded H2-D$^d$ or gTAX-loaded HLA-A*02:01, in the presence or absence of free peptides P18-I10 or motif (for H2-D$^d$) or HTLV-1 TAX (for HLA-A*02:01). The gating strategy for one of the sorting experiments is shown. In this example, yeast were incubated with 0.5 μg/ml H2-D$^d$ tetramer, and yeast expressing TAPBPR variants with the highest levels of binding were collected (magenta gate). MHC-I tetramer staining was below saturation (staining remained unsaturated up to at least 10 μg/ml tetramer). **(C-G)** Yeast-displayed TAPBPR libraries were sorted for binding to MHC-I tetramers and the enrichment or depletion of mutations were calculated following Illumina sequencing of the naive and sorted populations. Comparisons are shown between different sorting experiments for the enrichment ratios of nonsynonymous (black) and nonsense (red) mutations in the TAPBPR 24-35 loop. Data are highly correlated between replicate experiments using different concentrations of H2-D$^d$ (C) or HLA-A*02:01 (F). Enrichment ratios are qualitatively similar when free peptides P18-I10 (D), motif (E), or TAX (G) are co-incubated with H2-D$^d$ (D, E) or HLA-A*02:01 (G), except for an increase in stringency for the tightest affinity TAPBPR variants (indicated by an upwards trend in positive $\log_2$ enrichment ratios).

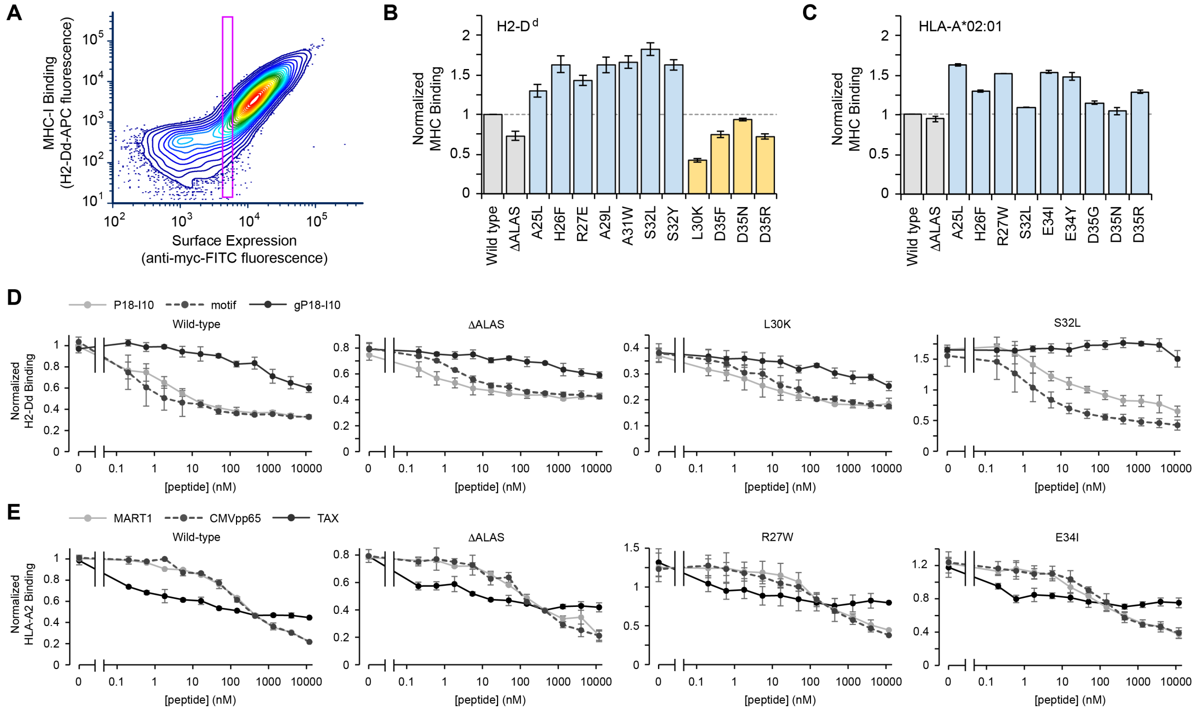


**Figure S5. TAPBPR mutants in the 24-35 loop with differences in binding to folded MHC-I. (A)** Yeast displaying surface TAPBPR were dual stained with P18-I10-loaded H2-$D^d$-APC tetramers and anti-myc-FITC for simultaneous detection by flow cytometry of bound MHC-I versus TAPBPR expression. Due to the use of avid MHC-I tetramers, binding to TAPBPR was found to be highly dependent on surface expression levels. To control for this, yeast were gated (magenta box) for low TAPBPR expression to minimize avidity effects and control for any differences in expression among TAPBPR mutants. Bound MHC-I (based on mean APC fluorescence) was measured within the magenta gate. **(B, C)** Validation by targeted mutagenesis of TAPBPR mutants predicted from the deep mutational scans to have reduced (orange) or increased (blue) binding to H2-$D^d$ (B; 1.0 µg/ml, mean ± SD from n = 4) or HLA-A*02:01 (C; 2.0 µg/ml, mean ± range from n = 2). TAPBPR ΔALAS (grey) has slightly reduced MHC-I binding. Mean fluorescence for MHC-I binding is normalized to wild type TAPBPR (grey). **(D, E)** Binding of H2-$D^d$ (D; 2.0 µg/ml, mean ± SD from n = 4) or HLA-A*02:01 (E; 2.0 µg/ml, mean ± range from n = 2) to TAPBPR-expressing yeast is competed by peptides. Shown are data from yeast expressing wild type TAPBPR, TAPBPR ΔALAS, and representative mutants. gP18-I10 is a low affinity peptide for H2-$D^d$ missing an N-terminal anchoring residue for the A pocket. Peptide sequences are: RGPGRAFVTI (P18-I10), GPGRAFVTI (gP18-I10), AGPARAAAL (motif), LLFGYPVYV (HTLV-1 TAX), ELAGIGILTV (MART1), NLVPMVATV (CMVpp65).

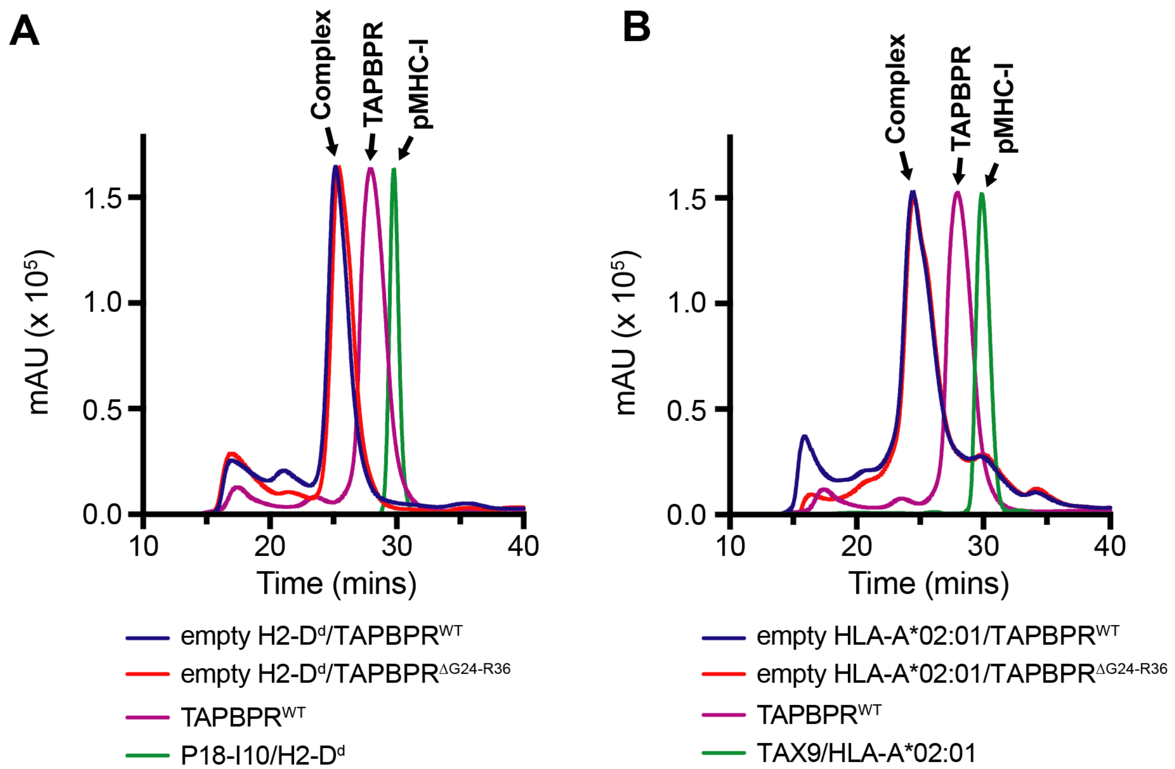


**Figure S6. Purification of peptide-deficient MHC-I/TAPBPR complexes.**
Size exclusion chromatographs (0.5 mL/min flow rate) of peptide-deficient MHC-I/TAPBPR complexes, relative to the free pMHC-I and TAPBPR species. photoP18-I10/H2-$D^d$/h$\beta_2$m **(A)** or photoFluM1/HLA-A*02:01/h$\beta_2$m **(B)** were mixed with TAPBPR at 1:1 molar ratio and incubated for 1 hour at 25°C. The mixture was then UV-irradiated at 365 nm for 1 hour followed by centrifugation at 13,000 r.p.m. for 10 min to remove precipitates. Pure 1:1 stoichiometric H2-$D^d$/h$\beta_2$m/TAPBPR (A) or HLA-A*02:01/h$\beta_2$m/TAPBPR (B) complexes were isolated by SEC with a Superdex 200 Increase 10/300 GL column at flow rate of 0.5 mL/min in 50 mM NaCl, 20 mM sodium phosphate pH 7.2. The exact procedure was followed for complexes containing either TAPBPR$^{WT}$ or TAPBPR$^{\Delta G24\text{-}R36}$, with chromatograms shown as different colors. Validation of peptide removal from the peaks at approx. 26 min corresponding to the MHC-I/TAPBPR complexes was achieved using liquid chromatography-mass spectroscopy (LC-MS) and carried out with passage through a Higgins PROTO300 C4 column (5 µm, 100 mm × 2.1 mm) followed by electron ion spray mass spectroscopy performed on a Thermo Finnigan LC-MS/MS (LTQ).

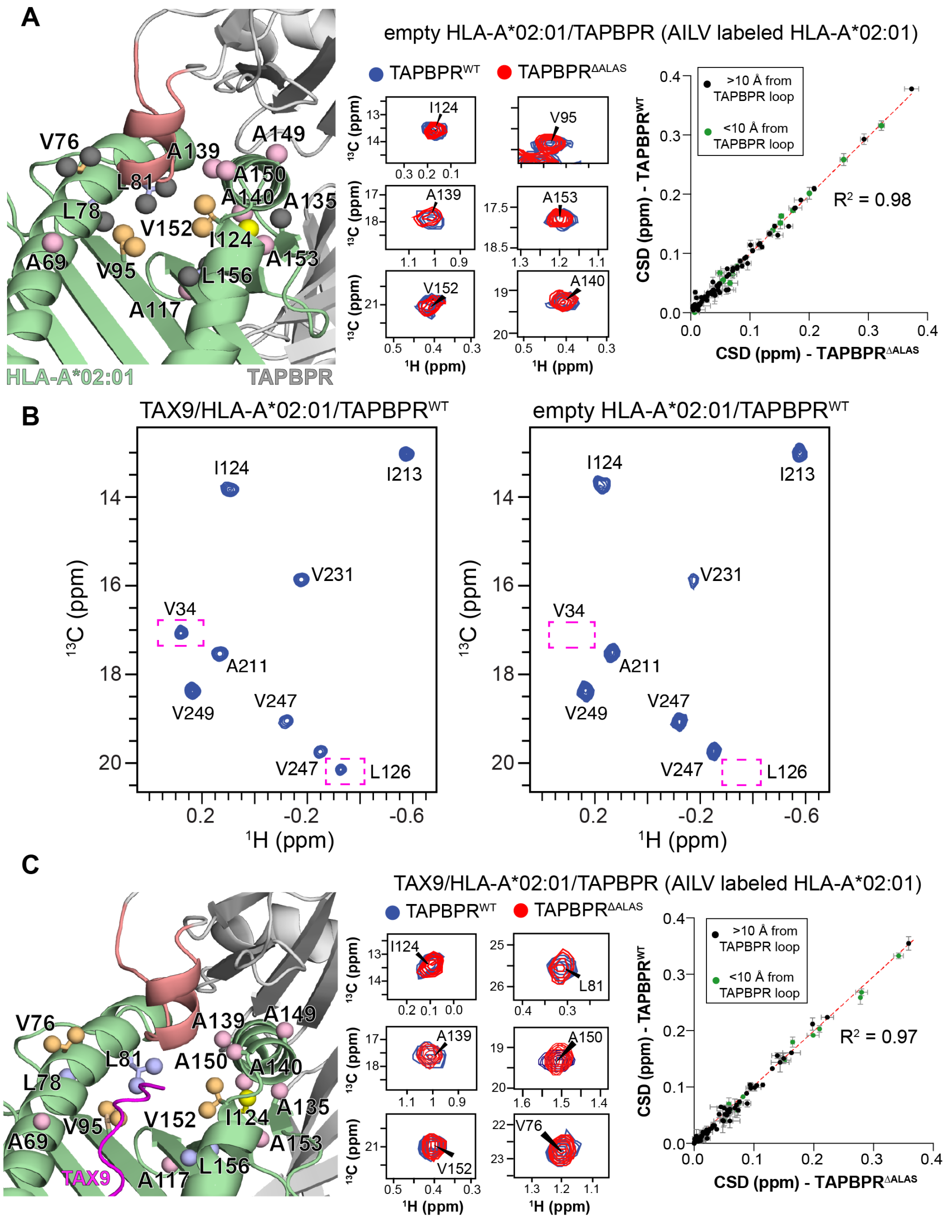


**Figure S7. The TAPBPR G24-R36 loop does not form a stable interaction with the HLA-A*02:01 groove. (A)** (Left) View of the peptide-deficient HLA-A*02:01/TAPBPR model (template PDB ID 5OPI) showing AILV methyl probes on HLA-A*02:01 (as spheres) within 10 Å angstroms of the TAPBPR G24-R36 loop (salmon). Methyl resonances of HLA-A*02:01 residues shown in black are missing in 2D $^{1}$H-$^{13}$C methyl HMQC spectra of peptide-deficient HLA-A*02:01/hβ$_2$m/TAPBPR complex due to conformational exchange induced line broadening. (Middle) Zoom in of representative peaks from 2D $^{1}$H-$^{13}$C methyl HMQC spectra of 80 μM peptide-deficient HLA-A*02:01 (AILV labeled)/hβ$_2$m in complex with TAPBPR$^{WT}$ (red) and TAPBPR$^{\Delta ALAS}$ (blue) recorded at 25°C at a $^{1}$H field of 800 MHz. (Right) Comparison of chemical shift deviation (CSD) values measured between free TAX9/HLA-A*02:01/hβ$_2$m and peptide-deficient HLA-A*02:01/hβ$_2$m in complex with TAPBPR$^{WT}$ or TAPBPR$^{\Delta ALAS}$. CSD values for HLA-A*02:01 methyl probes within 10 Å of the TAPBPR G24-R36 loop are shown in green, while those further than 10 Å are shown in black. The dotted red line is a linear fit of the data with

$R^2$ value noted. **(B)** Comparison of 2D $^1H$-$^{13}C$ HMQC spectra of 80 μM TAX9/HLA-A*02:01 (AILV labeled)/h$\beta_2$m/TAPBPR$^{WT}$ in the presence of 200 μM excess TAPBPR$^{WT}$ (left) with 80 μM peptide-deficient HLA-A*02:01 (AILV labeled)/h$\beta_2$m/TAPBPR$^{WT}$ (right). Dotted pink boxes highlight methyl resonances that become exchange broadened in the empty complex. **C.** (Left) View of the TAX9/HLA-A*02:01/TAPBPR model (template PDB ID 5OPI) showing AILV methyl probes on HLA-A*02:01 (as spheres) within 10 Å angstroms of the TAPBPR G24-R36 loop (salmon). (Middle) Zoom in of representative peaks from 2D $^1H$-$^{13}C$ methyl HMQC spectra of 80 μM TAX9/HLA-A*02:01 (AILV labeled)/h$\beta_2$m in complex with TAPBPR$^{WT}$ (red) and TAPBPR$^{\Delta ALAS}$ (blue) recorded at 25°C at a $^1H$ field of 800 MHz. (Right) Comparison of chemical shift deviation (CSD) values measured between free TAX9/HLA-A*02:01/h$\beta_2$m and TAX9/HLA-A*02:01/h$\beta_2$m in complex with TAPBPR$^{WT}$ or TAPBPR$^{\Delta ALAS}$. CSD values for HLA-A*02:01 methyl probes within 10 Å of the TAPBPR G24-R36 loop are shown in green, while those further than 10 Å are shown in black. The dotted red line is a linear fit of the data with $R^2$ value noted.

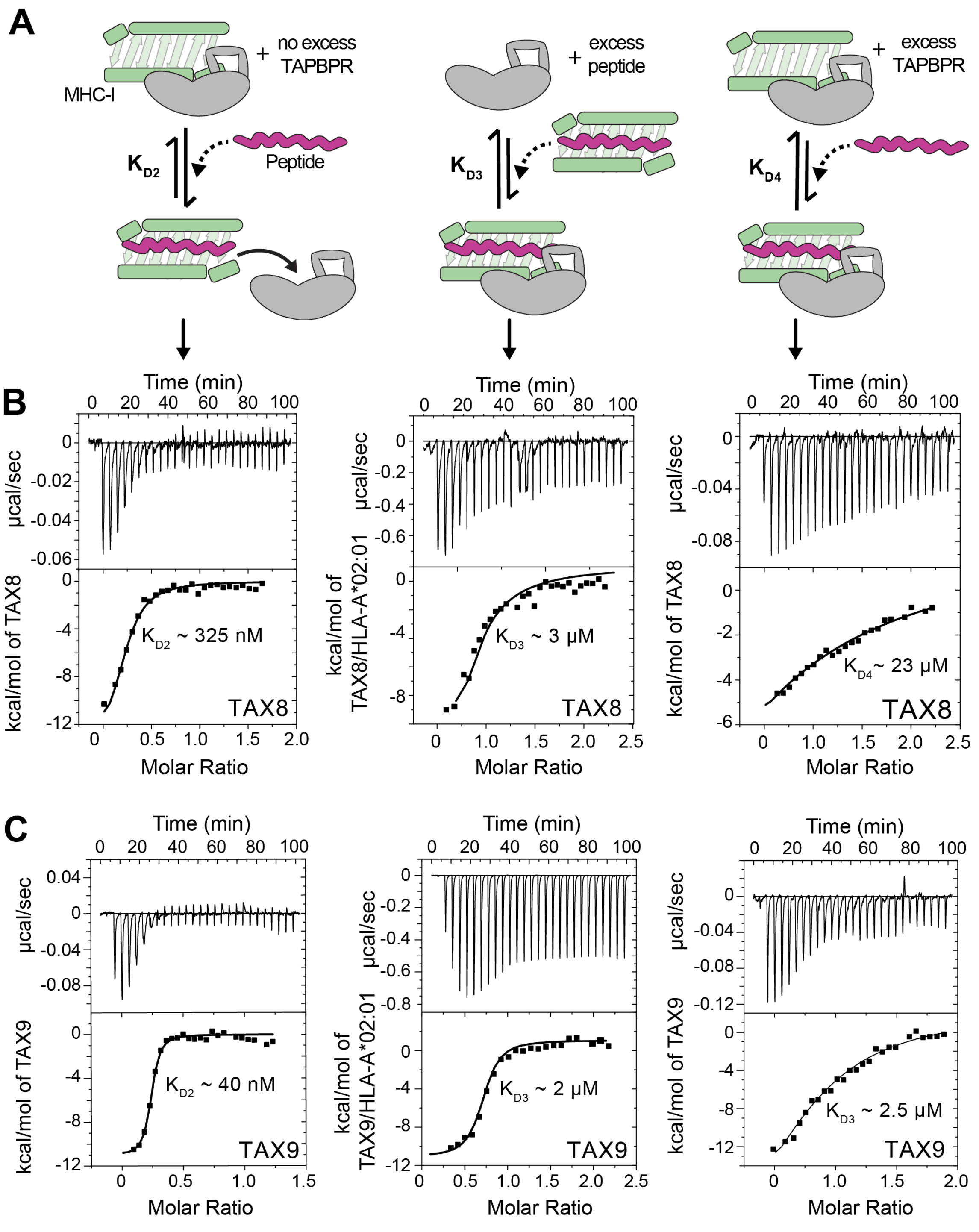


**Figure S8. Design of ITC experiments to examine specific steps of the thermodynamic cycle of TAPBPR mediated peptide exchange.** **(A)** Schematic of conditions in the calorimetry cell (top) and the syringe (titration represented by the dotted arrow) during ITC experiments for each step of the peptide exchange cycle. **(B)** and **(C)** Representative examples of raw ITC data for each step of the peptide exchange cycle for TAX8 and TAX9 peptide. The line represents the best fit of the data using a 1:1 model in Origin. Fitted apparent $K_D$ values are reported. Details of the experiments are outlined in *Materials and Methods*.

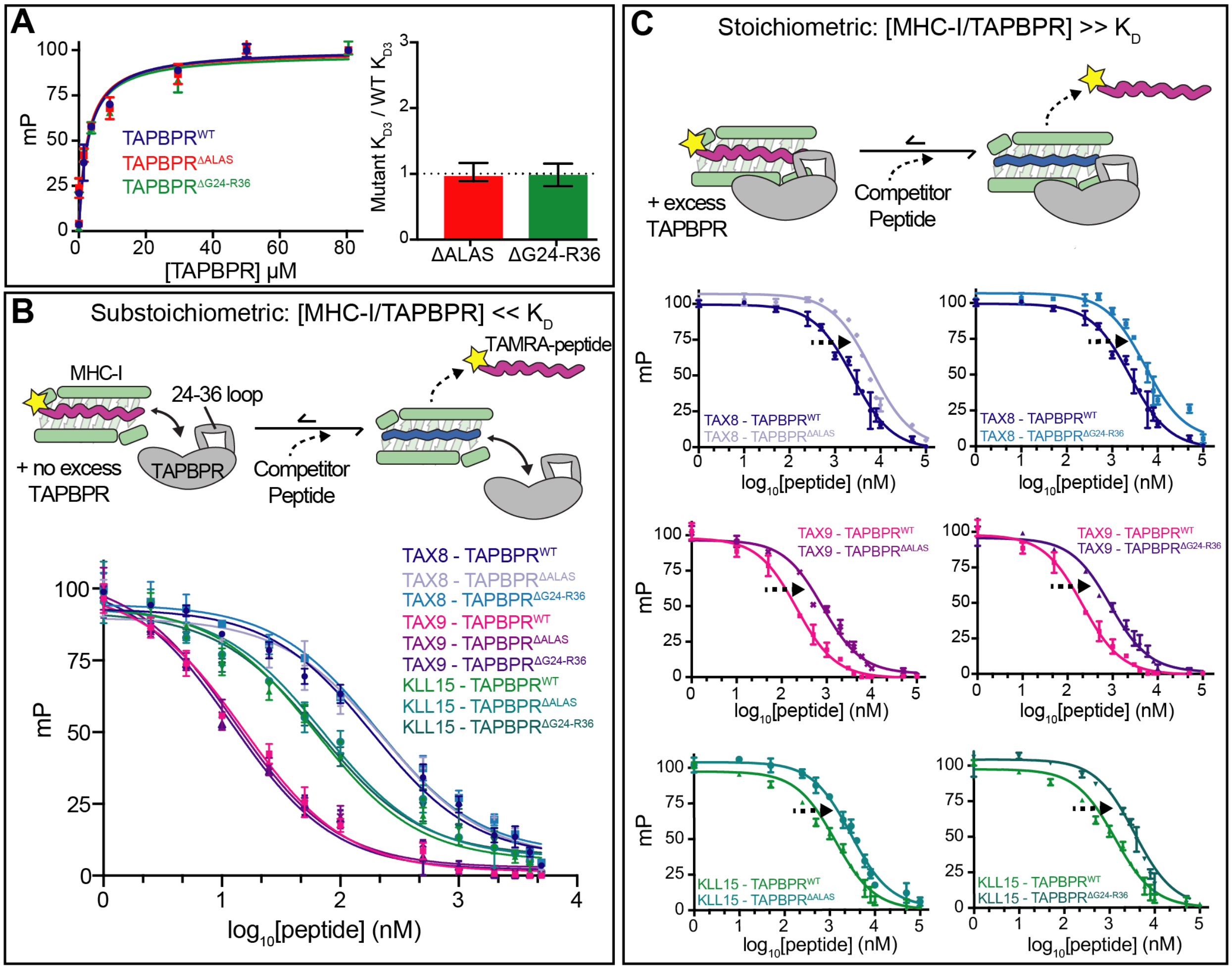


**Figure S9. Design of FP assays to examine specific steps of the thermodynamic cycle of TAPBPR mediated peptide exchange.** **(A)** FP experiments to probe the $K_{D3}$ step of the peptide exchange cycle. (Left) Titration of graded concentrations of $TAPBPR^{WT}$, $TAPBPR^{\Delta ALAS}$ or $TAPBPR^{\Delta G24\text{-}R36}$ into 1 nM TAMRA-TAX9 and 50 nM TAX8/HLA-A*02:01/h$\beta_2$m. (Right) Ratio of FP-determined $K_{D3}$ values for mutant / wild-type TAPBPR. **(B)** Schematic of the design of FP experiments performed under conditions of low TAPBPR binding to MHC-I. 50 nM of wild-type or G24-R36 loop mutant peptide-deficient HLA-A*02:01/h$\beta_2$m/TAPBPR complex is incubated with 1 nM TAMRA-TAX9 peptide and a range of concentrations of competitor peptide. millipolarization (mP) values are plotted as a function of the $\log_{10}$ peptide concentration for TAX8, TAX9 and KLL15 peptides. **(C)** Schematic of the design of FP experiments performed under 20-fold molar excess of TAPBPR. 50 nM of wild-type or G24-R36 loop mutant peptide-deficient HLA-A*02:01/h$\beta_2$m/TAPBPR complex is incubated with 1 nM TAMRA-TAX9 peptide, 2 μM TAPBPR (or $TAPBPR^{\Delta ALAS}$ or $TAPBPR^{\Delta G4\text{-}R36}$) and a range of concentrations of competitor peptide. mP values are plotted as a function of the $\log_{10}$ peptide concentration for TAX8, TAX9 and KLL15 peptides. Error bars were obtained from three technical replicates.

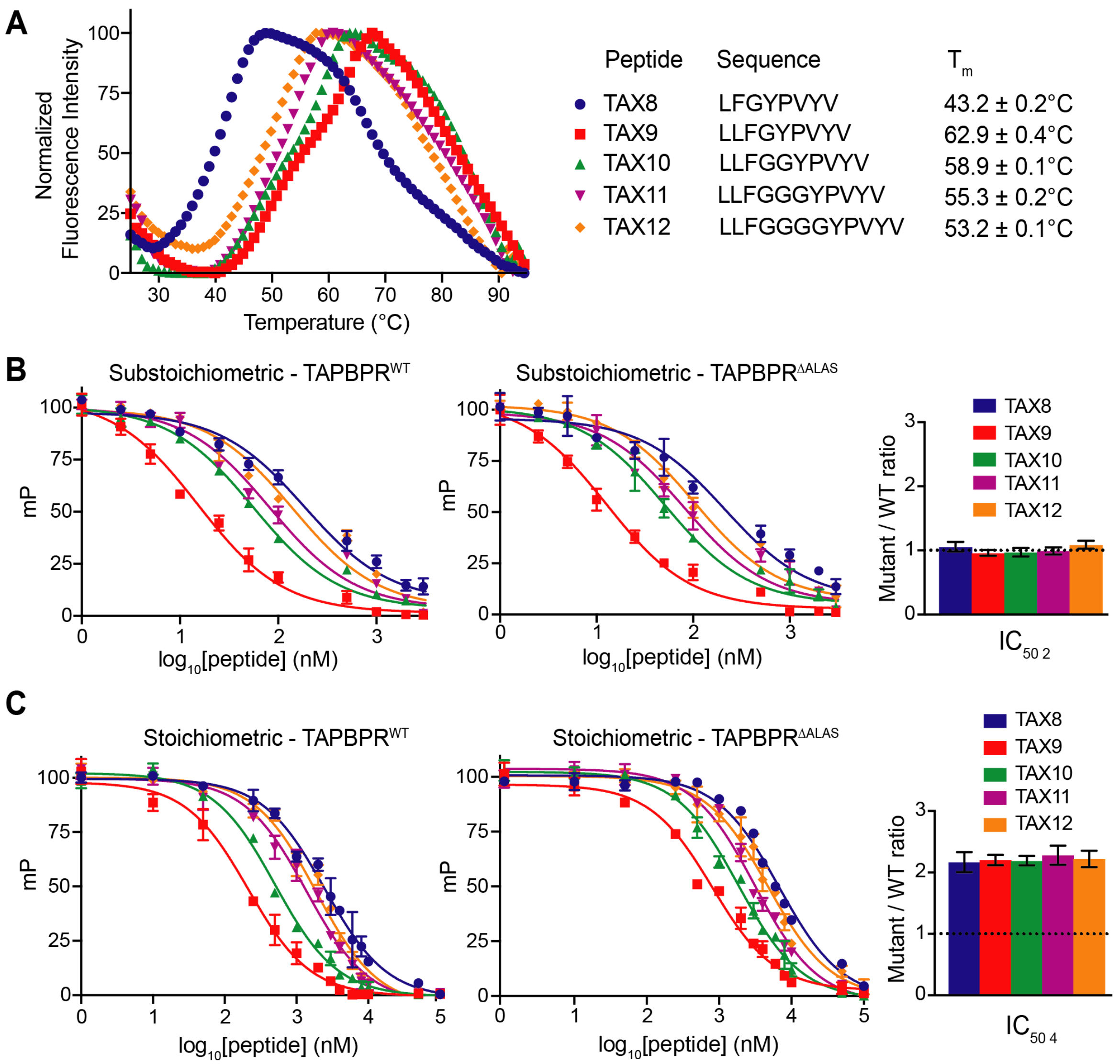


**Figure S10. Evaluation of peptide length dependence for TAPBPR mediated exchange.** (**A**) DSF experiments performed on 7 μM HLA-A*02:01/h$\beta_2$m in complex with different TAX length variants. The peptide sequence and determined $T_m$ values are shown. Errors were determined from three replicates. Fluorescence intensities were normalized for comparison. **(B)** (Left, Middle) FP performed under low TAPBPR binding conditions, 50 nM of peptide-deficient HLA-A*02:01/h$\beta_2$m/TAPBPR$^{WT}$ or HLA-A*02:01/h$\beta_2$m/TAPBPR$^{\Delta ALAS}$ complex is incubated with 1 nM TAMRA-TAX9 peptide and a range of concentrations of competitor peptide. millipolarization (mP) values are plotted as a function of the $\log_{10}$ peptide concentration for the TAX length variant peptides. (Right) Comparison of the ratio of FP determined $^2IC_{50}$ values for TAPBPR$^{\Delta ALAS}$ versus TAPBPR$^{WT}$. The dotted line represents no effect. **(C)** (Left, Middle) FP performed under 20-fold molar excess of TAPBPR, 50 nM of peptide-deficient HLA-A*02:01/h$\beta_2$m/TAPBPR$^{WT}$ or HLA-A*02:01/h$\beta_2$m/TAPBPR$^{\Delta ALAS}$ complex is incubated with 1 nM TAMRA-TAX9 peptide, 2 μM TAPBPR (or TAPBPR$^{\Delta ALAS}$) and a range of concentrations of competitor peptide. mP values are plotted as a function of the $\log_{10}$ peptide concentration for the TAX length variant peptides. (Right) Comparison of the ratio of FP determined $^4IC_{50}$ values for TAPBPR$^{\Delta ALAS}$ versus TAPBPR$^{WT}$. The dotted line represents no effect.

### Supplemental References


1. A. C. McShan, *et al.*, Molecular determinants of chaperone interactions on MHC-I for folding and antigen repertoire selection. *Proc. Natl. Acad. Sci. U. S. A.* **116**, 25602–25613 (2019).

2. E. Procko, *et al.*, Computational design of a protein-based enzyme inhibitor. *J. Mol. Biol.* **425**, 3563–3575 (2013).

3. S. J. Fleishman, *et al.*, Computational Design of Proteins Targeting the Conserved Stem Region of Influenza Hemagglutinin. *Science* **332**, 816–821 (2011).

4. P. Mali, *et al.*, RNA-Guided Human Genome Engineering via Cas9. *Science* **339**, 823–826 (2013).

5. M. Güell, L. Yang, G. M. Church, Genome editing assessment using CRISPR Genome Analyzer (CRISPR-GA). *Bioinformatics* **30**, 2968–2970 (2014).

6. J. Park, *et al.*, Structural architecture of a dimeric class C GPCR based on co-trafficking of sweet taste receptor subunits. *J. Biol. Chem.* **294**, 4759–4774 (2019).

7. D. M. Fowler, C. L. Araya, W. Gerard, S. Fields, Enrich: software for analysis of protein function by enrichment and depletion of variants. *Bioinforma. Oxf. Engl.* **27**, 3430–3431 (2011).

8. D. N. Garboczi, D. T. Hung, D. C. Wiley, HLA-A2-peptide complexes: refolding and crystallization of molecules expressed in Escherichia coli and complexed with single antigenic peptides. *Proc. Natl. Acad. Sci.* **89**, 3429–3433 (1992).

9. B. Rodenko, *et al.*, Generation of peptide-MHC class I complexes through UV-mediated ligand exchange. *Nat. Protoc.* **1**, 1120–1132 (2006).

10. G. I. Morozov, *et al.*, Interaction of TAPBPR, a tapasin homolog, with MHC-I molecules promotes peptide editing. *Proc. Natl. Acad. Sci. U. S. A.* **113**, E1006-1015 (2016).

11. C. Wang, P. Bradley, D. Baker, Protein-protein docking with backbone flexibility. *J. Mol. Biol.* **373**, 503–519 (2007).

12. D. J. Mandell, E. A. Coutsias, T. Kortemme, Sub-angstrom accuracy in protein loop reconstruction by robotics-inspired conformational sampling. *Nat. Methods* **6**, 551–552 (2009).

13. Y. Song, *et al.*, High-Resolution Comparative Modeling with RosettaCM. *Structure* **21**, 1735–1742 (2013).

14. A. C. McShan, *et al.*, Peptide exchange on MHC-I by TAPBPR is driven by a negative allostery release cycle. *Nat. Chem. Biol.* **14**, 811–820 (2018).

15. P. Rossi, Y. Xia, N. Khanra, G. Veglia, C. G. Kalodimos, 15N and 13C- SOFAST-HMQC editing enhances 3D-NOESY sensitivity in highly deuterated, selectively [1H,13C]-labeled proteins. *J. Biomol. NMR* **66**, 259–271 (2016).

16. F. Delaglio, *et al.*, NMRPipe: a multidimensional spectral processing system based on UNIX pipes. *J. Biomol. NMR* **6**, 277–293 (1995).

17. W. Lee, M. Tonelli, J. L. Markley, NMRFAM-SPARKY: enhanced software for biomolecular NMR spectroscopy. *Bioinforma. Oxf. Engl.* **31**, 1325–1327 (2015).
